# Nascent-Chain Interaction Networks and Their Effect on the Bacterial Ribosome

**DOI:** 10.1101/2022.10.31.514555

**Authors:** Meranda M. Masse, Valeria Guzman-Luna, Angela E. Varela, Rachel B. Hutchinson, Aniruddha Srivastava, Wanting Wei, Andrew M Fuchs, Silvia Cavagnero

**Author notes:** A.E.V.: School of Veterinary Medicine, University of Wisconsin-Madison, Madison, Wisconsin, 53706, USA. A.S.: School of Medicine and Public Health, University of Wisconsin-Madison, Madison, Wisconsin, 53705, USA. W.W.: AIDS Vaccine Research Laboratory, University of Wisconsin-Madison, Madison, Wisconsin, 53711, USA.

## Abstract

In order to become bioactive, proteins need to be biosynthesized and protected from aggregation during translation. The ribosome and molecular chaperones contribute to both of these tasks. While it is known that some ribosomal proteins (r-proteins) interact with ribosome-bound nascent chains (RNCs), specific interaction networks and their role within the ribosomal machinery remain poorly characterized and understood. Here, we find that RNCs of variable sequence and length (beyond the 1^st^ C-terminal reside) do not modify the apparent stability of the peptidyl-transferase center (PTC) and r-proteins. Thus, RNC/r-protein interaction networks close to the PTC have no effect on the apparent stability of ribosome-RNC complexes. Further, fluorescence anisotropy decay, chemical-crosslinking and Western blots show that RNCs of the foldable protein apoHmp_1-140_ have an N-terminal compact region (63–94 residues) and interact specifically with r-protein L23 but not with L24 or L29, at the ribosomal-tunnel exit. Longer RNCs bear a similar compact region and interact either with L23 alone or with L23 and another unidentified r-protein, or with molecular chaperones. The apparent strength of RNC/r-protein interactions does not depend on RNC sequence. Taken together, our findings show that RNCs encoding foldable protein sequences establish an expanding specific interaction network as they get longer, including L23, another r-protein and chaperones. Interestingly, the ribosome alone (i.e., in the absence of chaperones) provides indiscriminate support to RNCs bearing up to ca. 190 residues, regardless of nascent-chain sequence and foldability. In all, this study highlights the unbiased features of the ribosome as a powerful nascent-protein interactor.

**Significance Statement:** The presence of interactions between nascent chains bearing a foldable amino-acid sequence (with no signal or arrest tags) and specific ribosomal proteins has never been experimentally demonstrated, up to now. Here, we identify the ribosomal protein L23 as a specific nascent-chain interacting partner. We show that L23 establishes noncovalent contacts with nascent chains of the multi-domain foldable model protein apoHmp, which lacks signal/arrest sequences. Interactions with another ribosomal protein and with the trigger-factor and Hsp70 chaperones were also detected. Interestingly, ribosomal-protein/nascent-chain complexes have similar apparent stability, in the case of nascent chains of variable sequence and degree of foldability. These findings are significant because they advance our knowledge on ribosome-mediated nascent-protein interaction networks and suggest avenues to prevent undesirable aggregation.

## Introduction

Recent evidence suggests that the ribosome plays an active role in cotranslational protein folding (1–6). During translation, the nascent chain traverses the ribosomal exit tunnel, which is ca. 80-100 Å long, 10-35 Å wide (7–10) and is able to fit 30 to 40 nascent residues. (11–16) Within the ribosomal exit tunnel and its nearby regions across the highly negatively charged outer surface of the ribosome, (3) nascent chains encoding single-domain proteins become compact (17, 18) and acquire some secondary (19–23) and tertiary structure (5, 24–28). This set of observations proves the importance of the ribosome in nascent-protein structure formation.

During translation, the ribosome influences nascent protein chains at different levels. For instance, the inner geometry of the ribosomal exit tunnel favors formation of secondary nascent-chain structure, especially of α-helical (22, 29–31) or β-sheet nature (23). In addition, recent work showed that the ribosomal exit tunnel and vestibule enable acceleration of folding – but not unfolding – of a small single-domain protein, thereby stabilizing nascent protein chains relative to their free state in solution (32). On the other hand, the ribosome may also destabilize single-protein domains, in case the domain is far removed from the peptidyl transferase center (33). Further, the ribosome renders nascent chains soluble relative to the corresponding ribosome-released protein chains, thereby supporting cotranslational folding (34). Collectively, these results highlight the influence of the ribosome on nascent protein folding.

The ribosome is known to establish physical noncovalent interactions with nascent chains. As summarized in Table 1, these interactions have been identified in a variety of experimental studies and can be divided into three categories. Namely, *(i)* interactions between the ribosome and nascent chains carrying an N-terminal signal sequence (22, 35–39), *(ii)* interactions between the ribosome and nascent chains bearing C-terminal ribosome-stalling or arrest sequences (40–47), and *(iii)* interactions between nascent chains that lack N- or C-terminal ribosome-stalling or arrest sequences (48). Additional studies are consistent with the presence of ribosome-nascent-chain interactions, though they do not directly prove their existence (49–52).

**Table 1.** Summary of published experimental evidence on interactions between nascent protein chains (RNCs) and the ribosome based on single-particle cryo-EM or chemical crosslinking data.

| RNC | Technique | Interacting ribosomal protein | Reference |
| --- | --- | --- | --- |
| Proteins with an N-terminal signal sequence |  |  |  |
| Leader peptidase | photo-crosslinking | L4, L22 and L23 | 37 |
| Two regulatory ribosome stalling peptides | cryo-EM | L4 and L17 | 35 |
| Transmembrane segment of 111p membrane protein | photo-crosslinking | not determined | 16 |
| Signal anchor of FtsQprotein | photo-crosslinking | L23 and L29 | 39 |
| pOmt | photo-crosslinking | L23 and L24 | 36 |
| EsP1-25 | photo-crosslinking | L23 and L24 | 38 |
| Proteins with a C-terminal ribosome-stalling sequence |  |  |  |
| TnaC | photo-crosslinking | L22 and L24 | 41 |
| TnaC | cryo-EM | L22 | 43 |
| SecM | cryo-EM | L22 and L23 | 40 |
| SecM | mutagenesis | L22 | 42 |
| SecM | cryo-EM | L23 | 44 |
| Signal anchor of Dap2 Protein | photo-crosslinking | RpI4, Rp17 and RpI29 | 45 |
| SecM | NMR | not determined | 34 |
| Proteins with no N- or C-terminal tag |  |  |  |
| Phosphorylated insulin receptor (PIR) | chemical crosslinking | L23 | 48 |

While a body of research has explicitly addressed how the ribosome influences nascent polypeptides and proteins, little is known about how the nascent chain affects the properties of the ribosome. For instance, empty 70S ribosomes are known to be more prone to dissociation than ribosomes bearing both mRNA and peptidyl tRNA. This conclusion was reached upon addition of either ribosome-dissociation factors (53) or Hofmeister cosolutes (54–56). Other researchers established a similar finding upon depletion of magnesium ions (57–59). In a different study, addition of Hofmeister salts were employed to show that translation initiation complexes (including 70S in complex with initiator tRNA) disassemble more easily than peptidyl tRNAs bearing nascent chains (60). In this work, ribosomes carrying longer nascent chains were found to be less prone to dissociation (60). Other work has examined the effect of magnesium ions and other Hofmeister ions on empty 70S ribosome disassembly and how it changes sedimentation coefficients (61).

Yet, there is only a limited number of studies targeting the effect of non-Hofmeister denaturing agents on the ribosome. For instance, it is known that the 30S subunit disassembles in the presence of 6 molar urea (62). In addition, urea lowers the melting temperature and sedimentation coefficient of the 50S ribosomal subunit (63). The 30S subunit is more sensitive to thermal denaturation than the 50S subunit, and the 70S ribosome is most thermally stable (64). Most notably, 70S ribosomes bearing a nascent chain are less prone to chemical denaturation than empty ribosomes (33).

However, the effect of urea on the disassembly of various ribosome components (e.g. peptidyl transferase center -- a.k.a. PTC -- and ribosomal proteins) as a function of specific RNC characteristics (e.g., length and amino-acid sequence, interactions with ribosomal proteins) has not been characterized yet, to date. The present work moves initial steps towards filling this gap of knowledge.

We find that short peptidyl-tRNAs (snc-tRNAs) stabilize the 70S ribosome against denaturation by the non-Hofmeister cosolute urea, and we propose a multi-step model for the disassembly of ribosome-RNC complexes consistent with our findings. In addition, RNCs up to chain length 140 interact only with one ribosomal protein (r-protein), i.e., L23, in the vicinity of the tunnel exit. A wider interaction network, including one additional ribosomal protein and the trigger factor (TF) and Hsp70 chaperones, gets established as the nascent chain elongates further. Finally, our data also suggest that the interaction strength of individual RNC/r-protein populations does not vary significantly with RNC sequence and length. In all, our results show that (a) the interaction of foldable RNCs with the L23 ribosomal protein has been explicitly identified experimentally for the first time, and (b) the ribosome provides even, indiscriminate assistance to newly synthesized nascent protein chains, whether they are foldable or intrinsically disordered, via a powerful and highly specific interaction network.

## Results and Discussion

### The presence of very short nascent chains stabilizes the 70S ribosomal complex

We started our investigations by performing a series of sucrose-gradient studies on *E. coli* empty ribosomes and nascent-chain-loaded ribosomes. Our results, detailed in SI-Appendix Figure S1 and S3, showed that empty-70S ribosomes are more sensitive to urea denaturation than ribosomes bearing tRNAs linked to longer nascent chains. It appears that the aminoacyl-tRNA is responsible for most of the stabilizing effect (SI Appendix, Figs. S1 and S3). Interestingly, the length and amino-acid sequence of the nascent protein does not influence the urea sensitivity of ribosome-RNC complexes.

The sequence dependence of ribosome-nascent-chain interactions has been further explored in other parts of this work.

### Extending the nascent-protein-chain portfolio

The experiments described in the next sections employed a larger set of RNCs than the sucrose-gradient studies. Our specific purpose was to test the effect of RNC chain length, sequence and foldability on the apparent stability of the ribosome and to experimentally characterize the RNC-ribosome interaction network. First, we examined several constructs derived from the *Escherichia coli* protein flavohemoglobin (apoHmp). The structure and key building blocks of this protein analyzed in this work shown in Figure 1, a,b. The crystal structure of apoHmp comprises three domains, an N-terminal heme-binding (domain 1), a flavin adenine dinucleotide-binding (domain 2) and a C-terminal nicotinamide adenine dinucleotide-binding domain (domain 3), as schematically illustrated in Figure 1, b) (65). Hmp plays a key role in O_2_, NO and CO transport in *E. coli* and is involved in a variety of signaling pathways (66, 67). We also studied the behavior of the phosphorylated insulin receptor interacting region of the growth factor receptor-bound protein 14 (Grb14) from *Rattus norvegicus.* This protein, which is denoted here as PIR, is intrinsically disordered (68), i.e., an IDP. The specific nascent chain constructs of both proteins analyzed in this work are illustrated in SI-Appendix Figure S4.

**Figure 1.**
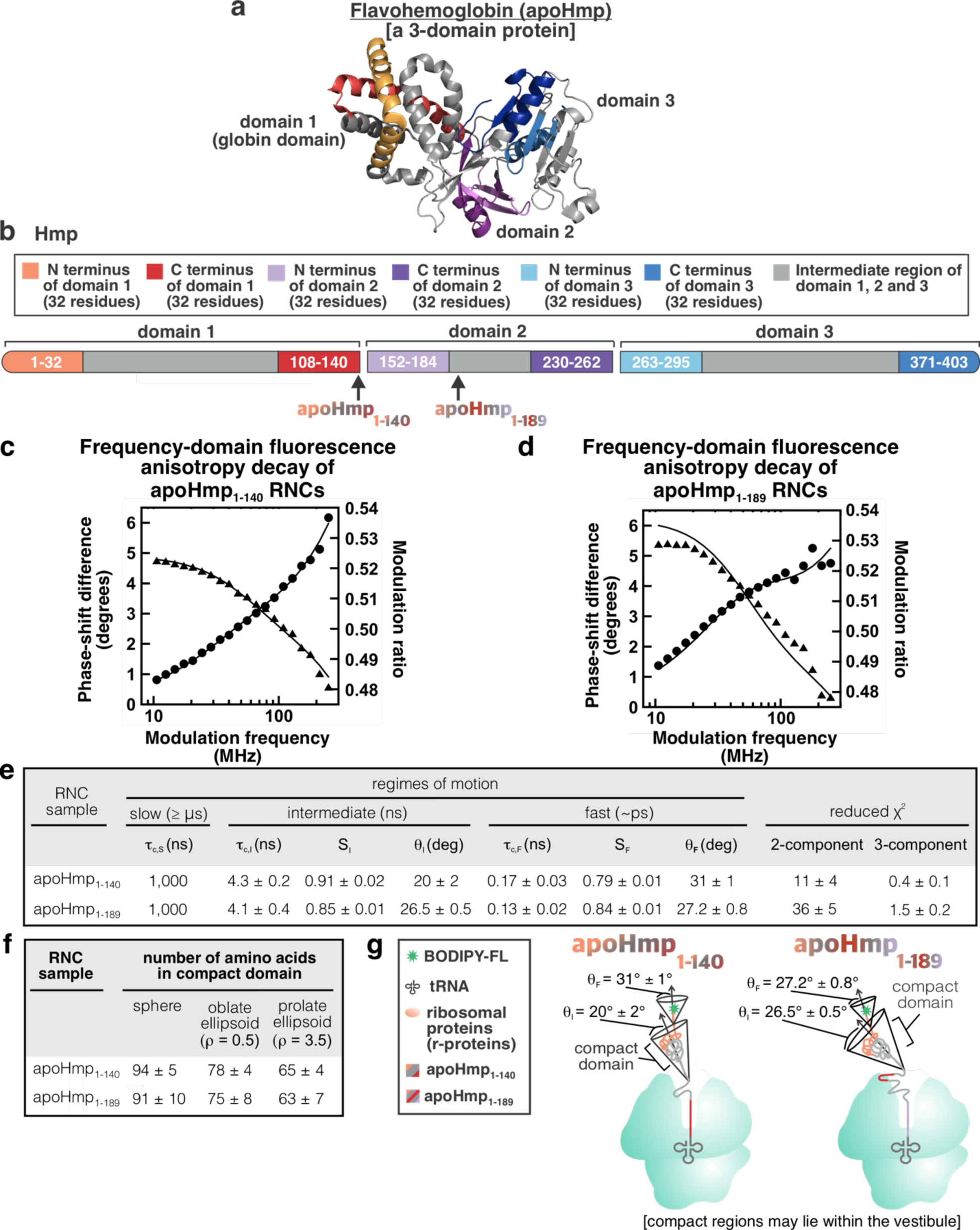
ApoHmp fluorescence-anisotropy decays reveal that apoHmpH nascent chains have a compact N-terminal region. **a)** *E. coli* flavohemoglobin (apoHmp) is the model foldable protein employed in this study. apoHmp has three domains (shown in red, purple and blue, PDB ID: 1GVH). **b)** Schematic representation of the three apoHmpH domains and their corresponding N and C termini. Representative frequency-domain fluorescence anisotropy decay data for **c)** apoHmp_1-140_ and **d)** apoHmp_1-189_ RNCs. **e)** Table summarizing anisotropy decay parameters including rotational correlation times (ρ_c_), order parameters (S) and cone semi-angles (θ). The S,I, and F subscripts denote slow-, intermediate-, and fast-timescale motions, respectively. Uncertainty values are reported as standard error for 3-5 independent experiments. Three-component fits were selected as best fits if they led to a ≥2.5-fold decrease in reduced χ^2^, relative to the two-component fit. The reduced χ^2^ of the chosen model is shown in bold. **f)** Table summarizing the number of amino acids comprising the RNC compact region, deduced from the τ_c,I_ rotational correlation time and assuming spherical, oblate ellipsoid, or prolate ellipsoid nascent-chain shapes. The parameter ρ denotes the axial ratio. **e)** Cartoon representation of apoHmp_1-140_ and apoHmp_1-189_ RNCs based on fluorescence anisotropy-decay data.

### Ribosome-bound apoHmp nascent chains of variable length are compact

Next, we performed fluorescence depolarization decay experiments in the frequency domain (69–71) to probe the rotational dynamics of nascent chains encoding foldable sequences. This technique has been previously employed to assess the rotational correlation time (ρ_c_) and amplitude of rotational motions of RNCs (17, 18, 34, 49, 72). The goal of this experiment was to probe whether RNCs harboring long nascent chains display any degree of compaction. We focused on RNCs of apoHmp_1-140_, corresponding to the N-terminal domain 1 of Hmp, and RNCs of apoHmp_1-189_, comprising Hmp domain 1 plus 49 C-terminal residues belonging to domain 2. Nascent proteins were site-specifically labeled at their N terminus with the BODIPY-FL fluorophore as described (17). Once information on nascent-chain compaction is in hand, the interplay between ribosome and nascent-chain sensitivity to urea denaturation can be more rationally explored and understood, as apparent in the sections below.

Representative data for apoHmp_1-140_ and apoHmp_1-189_ are shown in panels c and d of Figure 1, respectively. Both RNCs display informative frequency-domain anisotropy decay profiles. As shown in Figure 1e and consistent with the very low reduced χ^2^ values, the fits that include 3 rotational-tumbling components give the best results. Importantly, panels e and f of Figure 1 show that both apoHmp_1-140_ and apoHmp_1-189_ RNCs are characterized by an N-terminal compact domain. In both cases, this domain spans ca. 63 to 94 residues, depending on the exact shape. Note that RNC shape assessment is beyond the scope of this work. Regardless of the actual overall morphology of the compact domains, the fact that a similar-size compact domain is observed for both apoHmp_1-140_ and apoHmp_1-189_ suggests that both of these constructs undergo some partial folding on the ribosome. Surprisingly, the observed size of the compact domain of apoHmp_1-189_ RNCs is significantly smaller than domain 2, which comprises 140 residues (Fig. 1b). Therefore, biosynthesis of the additional 49 C-terminal amino acids is not sufficient to lead to complete folding of the N-terminal domain, for this protein.

In addition, the cone semi-angle analysis of the fluorescence anisotropy decay data (Fig. 1f) shows that the compact domain of Hmp_1-189_ RNCs spans a slightly wider cone semi-angle (26.5⁰ ± 0.5⁰ *vs* 20⁰ ± 0.2⁰) than Hmp_1-140_ RNCs, consistent with the fact that the latter construct is likely projecting further out from the ribosomal surface than the shorter Hmp_1-140_ construct.

In all, the fluorescence anisotropy data show that the Hmp_1-140_ and Hmp_1-189_ nascent chains are both comparably compact and no more than partially folded, while on the ribosome, with Hmp_1-189_ spanning a slightly wider cone semi-angle.

### The peptidyl transferase center site is largely unaffected by nascent-chain sequence and length, beyond 32 residues

Next, we probed whether nascent chains of different length, amino-acid sequence and foldability affect the apparent stability of specific regions of the ribosome. We directed our initial focus on the peptidyl transferase center (PTC) of the *E. coli* ribosome and we explored its urea sensitivity via a nascent-chain ribosome-release assay mediated by puromycin. Puromycin is a small-molecule antibiotic that induces premature release of nascent polypeptides from the ribosome. It mimics the adenosine-Phe portion of the CCA 3’ end of Phe-tRNA^Phe^ (Fig. 2a). Puromycin gets incorporated into nascent polypeptides as a result of nucleophilic attack of the carboxyl C -terminus of the nascent chain (73). On the other hand, this antibiotic works properly only if the A and P sites, which are entirely located within the 50S subunit, are intact (74). At high urea concentrations, the PTC is denatured and puromycin is no longer able to promote the release of nascent protein chains from the ribosome.

**Figure 2.**
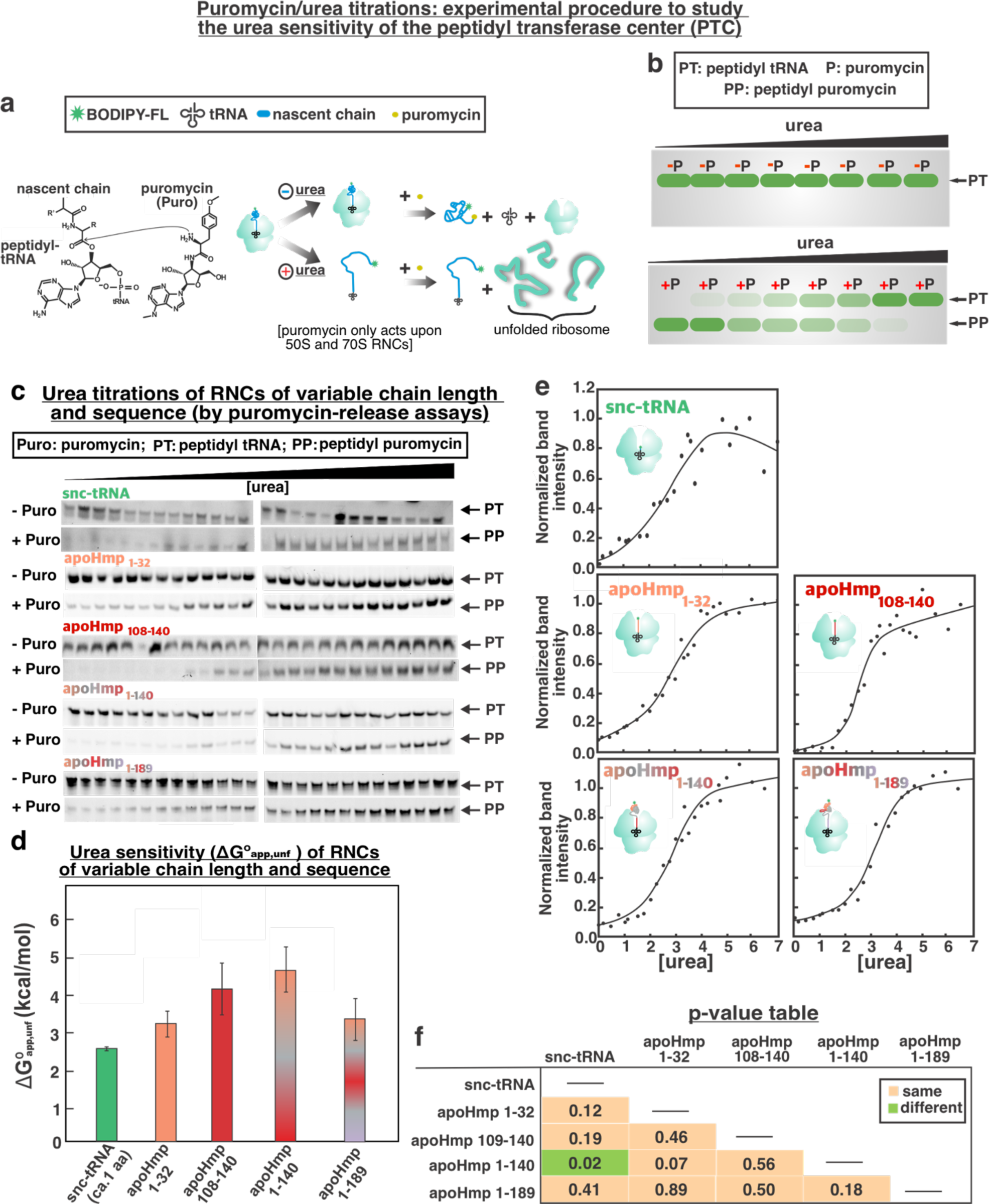
Sensitivity of peptidyl transferase center (PTC) of the ribosome to urea denaturation. **a)** Puromycin mimics the 3’ end of the amino acyl tRNA and covalently attaches to nascent chains, promoting their release from peptidyl tRNA. This process is accompanied by a significant decrease in molecular weight. **b)** Low-pH SDS-PAGE is employed to quantify peptidyl-tRNA (PT) band intensities as a function of urea concentration. Puromycin is unable to perform its function at high urea concentration because of pervasive PTC unfolding (SI Appendix, Fig. S1). **c)** Representative low-pH gels showing that peptidyl tRNA bands are unaffected by high urea concentrations. Addition of puromycin (1 mM) results in a decrease in peptidyl-tRNA (PT) band intensities, at low urea concentration. **d)** Block diagram mapping apparent unfolding free energy values (ΔG°_app,unfold_). Error bars denote standard errors based on 2-7 experiments. **e)** Representative urea titration curves corresponding to the raw data in panel c. Data were fit to a two-state unfolding expression (see Methods section for details). **f)** P*-*value table for a two-tailed Student’s test assuming unequal variances (Welch’s t-test). Green and orange boxes denote statistically different and statistically equivalent data, respectively, according to a ≥ 95% confidence interval.

The main features of the puromycin ribosome-release assay and its urea-concentration dependence are illustrated in Figure 2a, b. In short, RNC reactivity to puromycin is monitored via low-pH gels (75) as a function of increasing urea concentration.

We performed low-pH SDS-PAGE analysis on a variety of apoHmp RNCs comprising variable chain-length values (SI Appendix, Fig. S4), net charge and hydrophobicity, and degree of folding. The target nascent protein chains included snc (i.e., a 1-3 amino acid chain derived from apoHmp), apoHmp_1-32_, apoHmp_108-140_, apoHmp_1-140_ and apoHmp_1-189_. These constructs specifically enabled us to probe differences between nascent-chain sequences located within the ribosomal exit tunnel core, as well as partially and fully folded nascent-chain domains. Low-pH SDS-PAGE was used to generate peptidyl tRNA (PT) bands whose intensities were quantified as a function of increasing urea concentration. The peptidyl-puromycin band (PP) was not used to track ribosome release because of its highly environmentally sensitive fluorescence intensity. Titration curves were generated based on the relative band intensities of the PT bands of each construct, reporting on the extent of puromycin reactivity (Fig. 2c). The data were then fit to an equation relating the experimental observable at variable urea concentrations to the apparent stability (ΔG°_app, unfold_) of each construct, following the general procedures by Santoro and Bolen (76, 77) (see Methods).

As shown in Figure 2d-f, the results indicate that the apparent stability of the PTC center, monitored via puromycin activity assays, is very similar for all nascent-chain construct. Moreover, the two-tailed Student’s test assuming unequal variances (Welch’s test) shows that most constructs display equivalent behavior (Fig. 2f). In other words, most constructs contribute to a similar extent to the apparent stability of the PTC, amounting to ca. 2.5 – 4.5 kcal/mol. Therefore, the urea sensitivity of the PTC does not depend on nascent-chain properties. and may be dominated by the stabilizing effect of the tRNA. The results also suggest that the specific tRNA sequence does not have an effect either. An exception is provided by snc and apoHmp_1-140_ nascent proteins, given that the apoHmp_1-140_ construct displays a statistically significant PTC-stabilizing role. This feature is likely not due to the compact partially folded state of apoHmp_1-140_, given that fluorescence anisotropy shows that nascent apoHmp_1-189_ has a comparable degree of compaction.

Note that ribosome disassembly and unfolding due to denaturing agents is an irreversible process (78). Therefore, true thermodynamic ΔG°_unfold_ values cannot be obtained, upon treating the ribosome with denaturing agents. Thus, we refer to the ΔG°_unfold_ values derived in this and the following sections of this work as *apparent stability* values. The mere function of these quantities, which were derived from data fitting of urea-titrations, is to describe the urea sensitivity – and not the thermodynamic stability – of specific portions of *E. coli* ribosomes.

Overall, our puromycin titration data show that the nature of the nascent chain has a weak effect on the urea sensitivity of the PTC, in *E. coli*. Next, we explored the effect of nascent-chain properties on the apparent stability of ribosomal proteins.

### The global urea sensitivity of ribosomal proteins is largely unaffected by nascent-chain sequence and length

The global urea sensitivity of ribosomal proteins (r-proteins) was probed via urea titrations based on Trp fluorescence emission. Trp is a well-known fluorescent reporter, and its emission properties are highly environmentally sensitive. Upon inspection of the *E. coli* ribosomal-protein sequences via the 2WWL and 2WWQ Protein Data Bank (PDB) files, we ascertained that the r-proteins of the ribosome comprise a total of 32 Trps (Fig. 3a) roughly uniformly dispersed throughout the ribosomal structure. To gain insights at the highest possible resolution, we focused on all the apoHmp nascent chains listed in SI-Appendix Figure S4e, and on the PIR nascent chain, also listed in the same figure. Note that apoHmpH nascent chains only contribute two additional Trps at positions 120 and 149, and PIR only contributes on Trp at position 44. We regard these residues, present in some of the constructs, as contributing negligibly to the overall Trp fluorescence emission. In essence, the readout of this assay is dominated by the much larger contributions arising from the 32 Trps interspersed across the r-proteins. Urea titrations were carried out and Trp fluorescence emission was monitored (Fig. 3c). Spectral shifts were regarded as reporters of r-protein folding, and centers of mass of emission spectra were assessed to generate titration curves reporting on the urea sensitivity of r-proteins. Urea titration data were processed according to Santoro and Bolen (76, 77). Individual representative titration curves are shown in Figure 3d.

**Figure 3.**
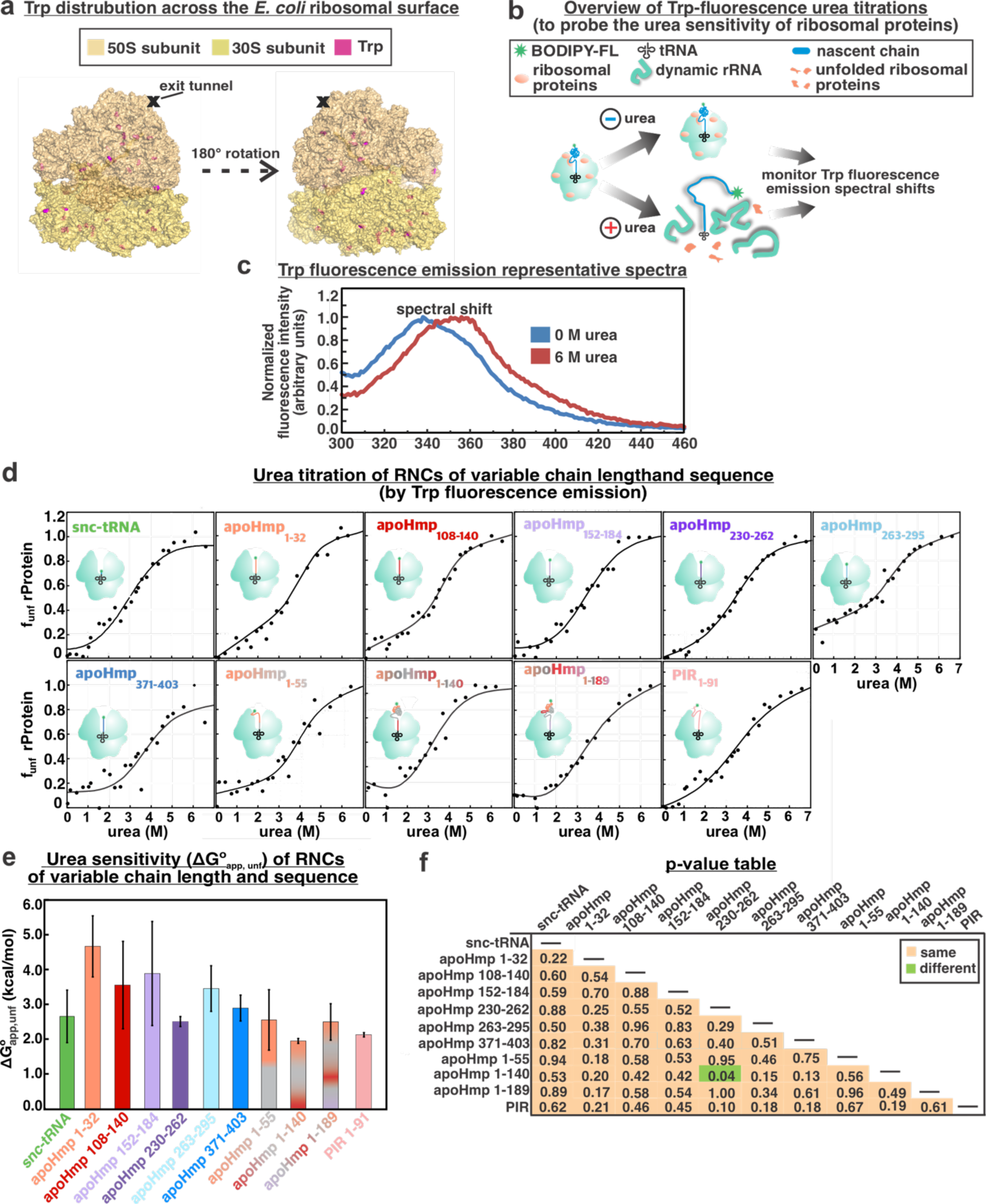
Sensitivity of ribosomal proteins to urea denaturation. **a)** The ribosomal proteins (r-proteins) of the *E. coli* 70S ribosome include a total of 32 Trp residues (magenta). Sixteen Trps are located within the proteins of the 50S subunit and sixteen are found in the r-proteins of the 30S subunit (PDB IDs: 2WWL and 2WWQ). **b)** Schematic illustration of the basic methodology followed to assess r-protein stability. **c)** The Trp fluorescence emission band becomes red-shifted at increasing urea concentrations due to environmental changes towards a more polar medium experienced by the Trp fluorophore. **d)** Representative urea titration curves obtained by steady-state fluorescence spectroscopy of snc-tRNA, apoHmp_1-32_, apoHmp_108-140_, apoHmp_152-184_, apoHmp_230-262_, apoHmp_263-295_, apoHmp_371-403_, apoHmp_1-55_, apoHmp_1-140_, apoHmp_1-189_ and PIR respectively. The symbol f_unf_ denotes change in the fraction of unfolded r-proteins, as shown in equation 4. **e)** Apparent unfolding free energy (ΔG°_app,unfold_) of each construct. Error bars denote the standard error based on 2-3 experiments. **f)** P*-*value table for a two-tailed Student’s T-test (Welch’s test) comparing the apparent free energy of unfolding values of each construct. Green and orange boxes denote statistically different and statistically equivalent data, respectively, according to a ≥ 95% confidence interval.

The ΔG°_app, unfold_ for each construct are plotted in Figure 3e and corresponding t-test values are tabulated in Figure 3f. The large majority of the constructs show statistically similar results, with ΔG°_app, unfold_ values ranging from 2 to 5 kcal/mol. Hence, the presence of peptidyl tRNA, regardless of nascent-chain characteristics, does not affect the urea sensitivity of r-proteins. As shown in sections below, some nascent chains interact with specific ribosomal proteins. On the other hand, these interactions are not sufficiently strong to be detected via this assay, which monitors the overall sensitivity to urea of all r-proteins.

### Sequence of events leading to ribosome disassembly: role of tRNA and nascent proteins

A proposed equilibrium ribosome disassembly mechanism based on the data of Figures 1 through 3 is shown in SI-Appendix Figure S6. The simple steps displayed in panel a of SI-Appendix Figure S6 pertain to empty 70S ribosomes and are consistent with the sucrose gradient data. The ribosome starts disassembling into its component 50S and 30S subunits at 1 M urea. More extensive subunit disassembly together with subunit unraveling follows at higher urea concentrations. This process is accompanied by pervasive r-protein and rRNA conformational heterogeneity, likely due to disruption of secondary and tertiary structure leading to r-RNA and r-protein unfolding. Panel b of the same figure shows how the process gets modified if the ribosome carries aminoacyl or peptidyl tRNA. Briefly, in this case the ribosomal-subunit disassembly occurs at higher urea concentrations (1 -2 M). Given the weak dependence of the sucrose-gradient and puromycin assays on the nature of the nascent chain, we deduce that the tRNA likely dominates the effect and that the length, hydrophobicity, net charge and foldability of apoHmp nascent chains do not play a stabilizing role in ribosome stability.

### Nascent chains of apoHmp_1-140_ and apoHmp_1-189_ interact with ribosomal protein L23

The data of Figure 3 showed that the apparent stability of r-proteins is not influenced by the presence of either intrinsically disordered or foldable nascent chains of variable length and physical properties. On the other hand, nascent chains encoding the intrinsically disordered protein PIR are known to interact with the r-protein L23 and L29 (48). It is reasonable to imagine that these interactions are not sufficiently strong to be detected by the Trp-fluorescence assay of Figure 3, which monitors Trp environment across the entire ribosome. Hence, the data in Figure 3 do not preclude the presence of RNC/ribosome interactions. Hence, it is compelling to explore whether these interaction exist. As shown in Figure 4a, the *E. coli* ribosome comprises several proteins within the region near the exit-tunnel.

**Figure 4.**
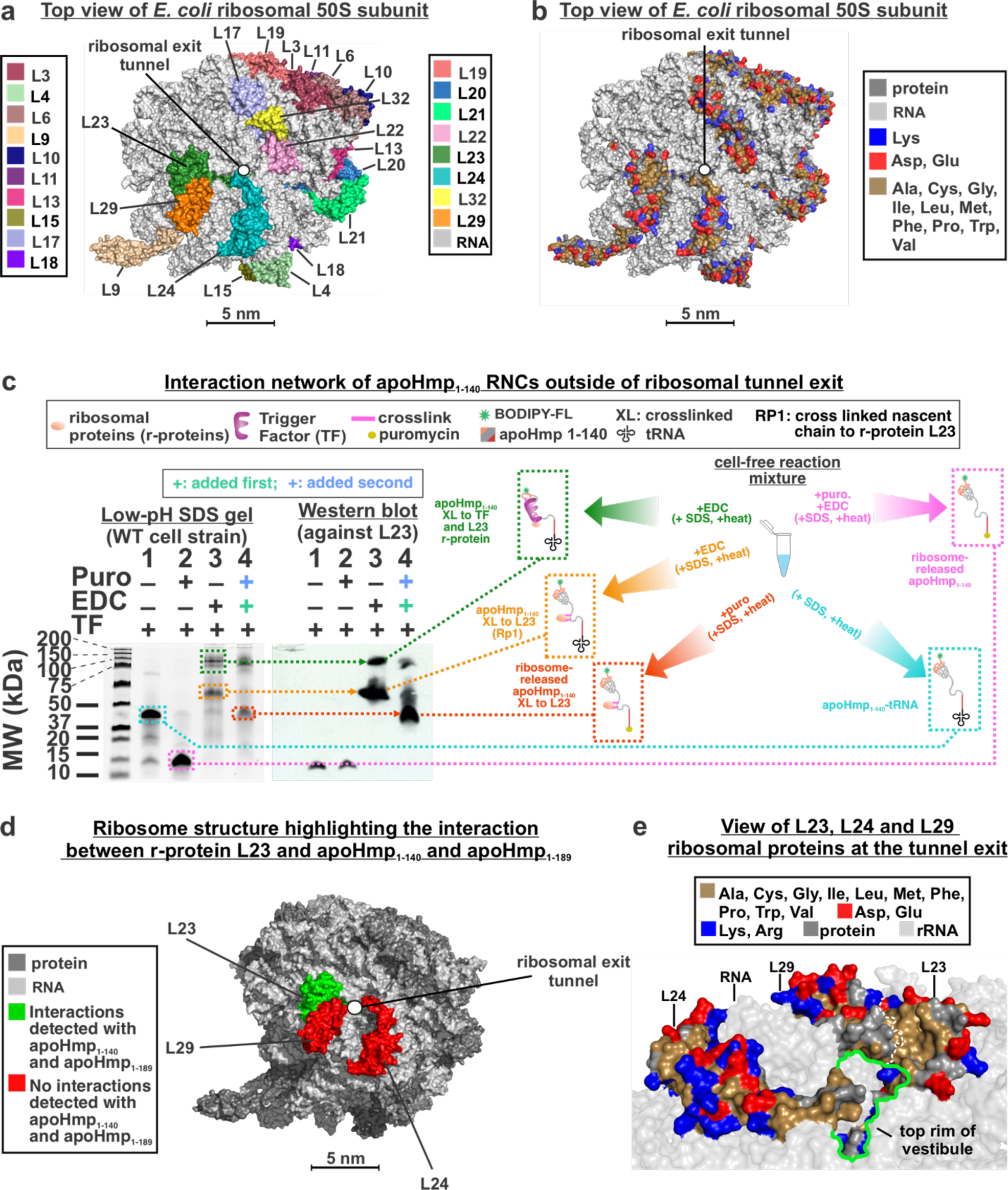
Crosslinking patterns of apoHmp_1-140_ RNCs and identification of interacting r-proteins. **a)** Top view of 50S subunit of the *E. coli* ribosome highlighting the ribosomal proteins (r-proteins). Figure has been modified from (48) under a Creative Commons Attribution 4.0 International license. **b)** Top view of 50S *E. coli* ribosome highlighting r-protein charged and nonpolar residues. Figure has been modified from (48) under a Creative Commons Attribution 4.0 International license. **c)** SDS-PAGE and Western blot data identifying r-proteins interacting with apoHmp_1-140_ RNCs in the absence and presence of the EDC crosslinker and the ribosome-release agent puromycin. Data show that the L23 r-protein interacts with apoHmp_1-140_ RNCs. Corresponding data employing antibodies against L24 and L29 r-proteins, showing no interactions, are available in the SI Appendix. **d)** Top view of *E. coli* 50S ribosomal subunit highlighting the r-proteins that either interact (green) or do not interact (red) with apoHmp_1-140_ and apoHmp_1-189_ RNCs. **e)** Side view of r-proteins near the vestibule of the ribosomal exit tunnel. Figure has been modified from (48) under a Creative Commons Attribution 4.0 International license.

In this work, we focused on the L23, L24 and L29 r-proteins, which are closest to the vestibule and outer region of the ribosomal exit tunnel. We explored the interaction patterns of the apoHmp_1-140_ and apoHmp_1-189_ nascent chains, which populate compact states while ribosome-bound (Fig. 1 c-g). The well-characterized zero-length chemical crosslinker carbodiimide 1-ethyl-3-[3-dimethylaminopropyl] carbodiimide hydrochloride (EDC) was employed (48, 79, 80). To carry out chemical crosslinking of nascent chains, we followed known procedures (48) involving a combination of low-pH SDS-PAGE (75) and Western blotting in the absence and presence of the trigger factor (TF) chaperone. Notably, EDC enables detecting existing noncovalent interactions, though it does not provide an accurate quantitation of interacting populations, as discussed at length by Guzman-Luna *et al.* (48). Yet, EDC is an extremely valuable tool to detect protein-protein interactions within the ribosome-nascent-chain complex. Site-specific fluorescence labeling of nascent proteins at their N terminus enables focusing exclusively on interactions involving the nascent protein. Low-pH-gel and Western-blot were collected to explore interactions between nascent chains and r-proteins. These interactions have not been directly detected before, in the case nascent chains lacking arrest or signal sequences. It is worth noting that EDC does not have high accessibility within the exit-tunnel core (48). Therefore, detecting interactions within the tunnel core, including corresponding Western-blotting work, would not have been informative within our experimental setup.

It is also important to mention that, under our experimental conditions, EDC does not report on interactions involving nascent protein chains and ribosomal RNA (rRNA). In the presence of imidazole, crosslinks between RNA 5’ phosphate and aliphatic amines of proteins is known to take place (79). However, our samples did not contain imidazole, and this chemical would anyways be unable to detect interactions not involving the 5’ end of RNA. Therefore, even in the presence of imidazole, EDC would likely underestimate all potential interactions with RNA. Thus, interactions between nascent proteins and r-RNA are beyond the scope of this study.

The data for apoHmp_1-140_ and apoHmp_1-189_, shown in Figures 4c and 5a,b, respectively, indicate that both nascent proteins interact with ribosomal protein L23. Western blotting was also carried out with antibodies against ribosomal proteins L24, L29 (SI Appendix, Figs. 7 and 9). However, no interactions between these two r-proteins and apoHmp_1-140_ and apoHmp_1-189_ were detected.

**Figure 5.**
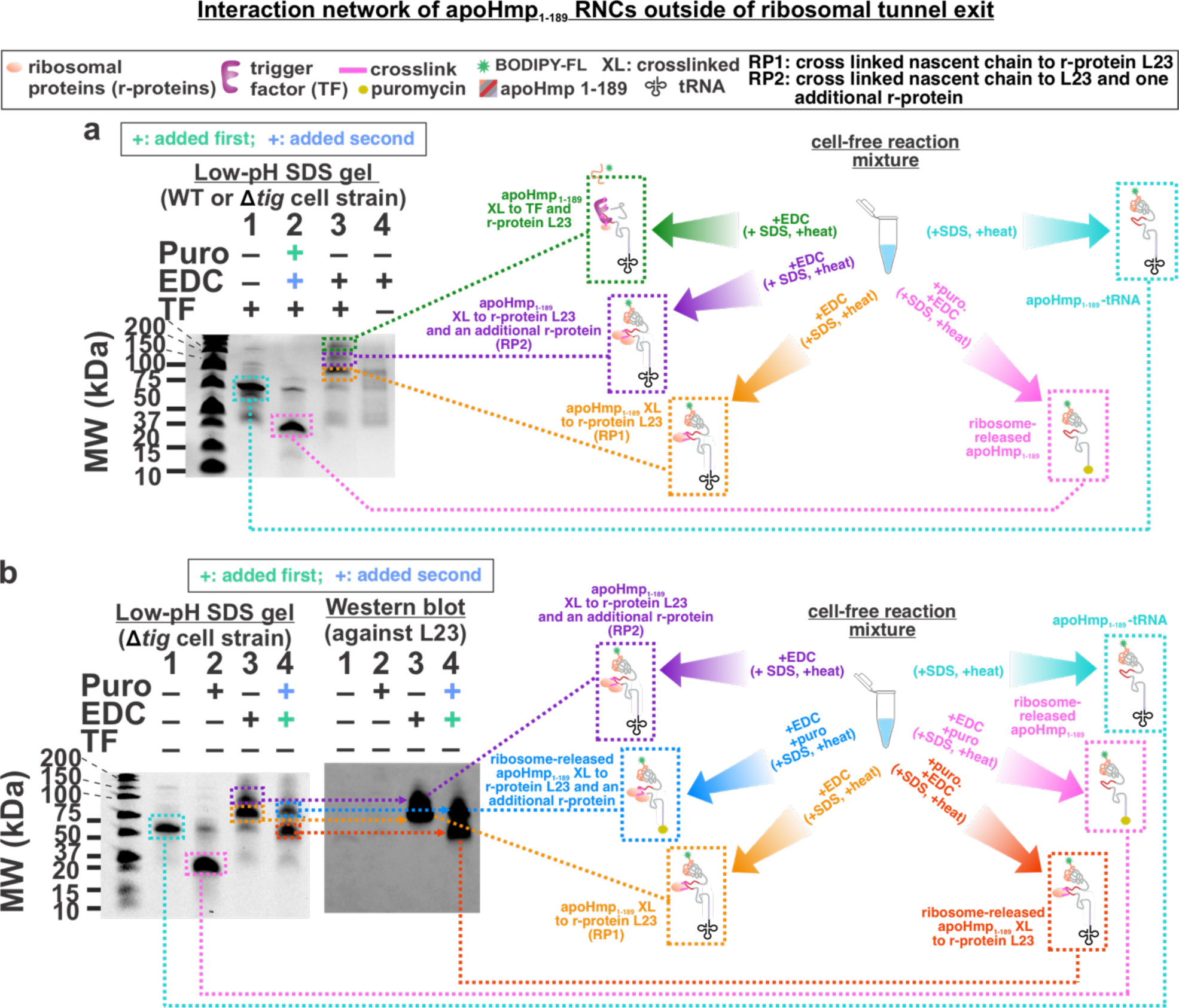
Crosslinking patterns of apoHmp_1-189_ RNCs and identification of interacting proteins. **a)** Low-pH SDS-PAGE analysis of apoHmp_1-189_ RNCs in the absence and presence of the EDC crosslinker, TF chaperone and the RNC ribosome-release agent puromycin. **b)** Side-by-side SDS-PAGE and Western blot data identifying the interaction network of apoHmp_1-189_ RNCs in the absence and presence of EDC, puromycin and TF chaperone. The L23 r-protein is found to interact with apoHmp_1-189_ RNCs. Corresponding data employing antibodies against L24 and L29 r-proteins, showing no interactions, are available in the SI Appendix.

**Figure 6.**
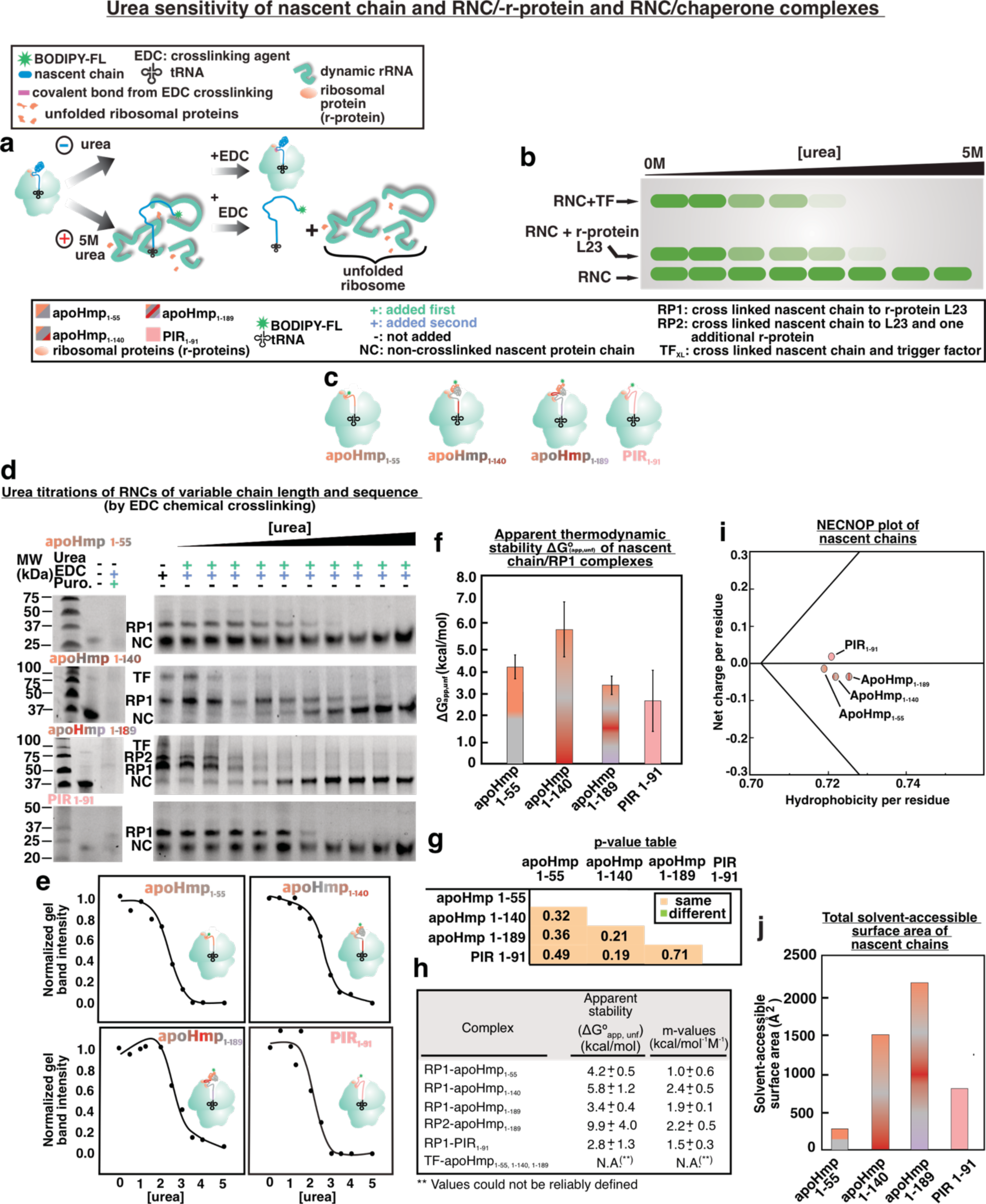
EDC-crosslinking-detected urea titrations showing the apparent stability of RNC/-r-protein complexes. **a)** Scheme showing the expected effect of urea addition on RNCs and **b)** corresponding low-pH SDS-PAGE gels., The gels were used to assess the extent of crosslinking, which is a reporter of RNC/r-protein apparent stability at different urea concentrations. **c)** Four RNCs were tested in these experiments:apoHmp_1-55_, apoHmp_1-140_, apoHmp_1-189_ or PIR_1-91_. Note that PIR_1-91_ is an intrinsically disordered protein (IDP). **d)** Representative SDS-PAGE analysis. The gel bands are reporters of the apparent stability of RNCs complexes with either L23 or TF. **e)** Representative urea titrations of apoHmp_1-55_, apoHmp_1-140_, apoHmp_1-189_ and PIR complexes. **f)** ΔG°_app,unfold_ values in the presence of low concentrations of chaperones (2-15 nM TF, and 0.5 µM, 0.04µM and 0.05µM DnaK, DnaJ and GrpE, respectively). Error bars denote standard errors based on 2-3 experiments. **g)** P-value table for a two-tailed Student’s T-test (Welch’s test), comparing apparent free energies of unfolding. Green and orange boxes denote statistically different and statistically equivalent data, respectively, according to a ≥ 95% confidence interval. **h)** Table displaying relevant ΔG°_app,unfold_ and m-values . **i)** NECNOP plot (96) displaying the net charge per residue as a function of hydrophobicity per residue of the PIR_1-91,_ apoHmp_1-55,_ apoHmp_1-140,_ apoHmp_1-189_ protein chains. **j)** total solvent-accessible surface area of each tested polypeptide, computed for hypothetical fully extended chains. Values were obtained via the Surfracer program (97).

**Figure 7.**
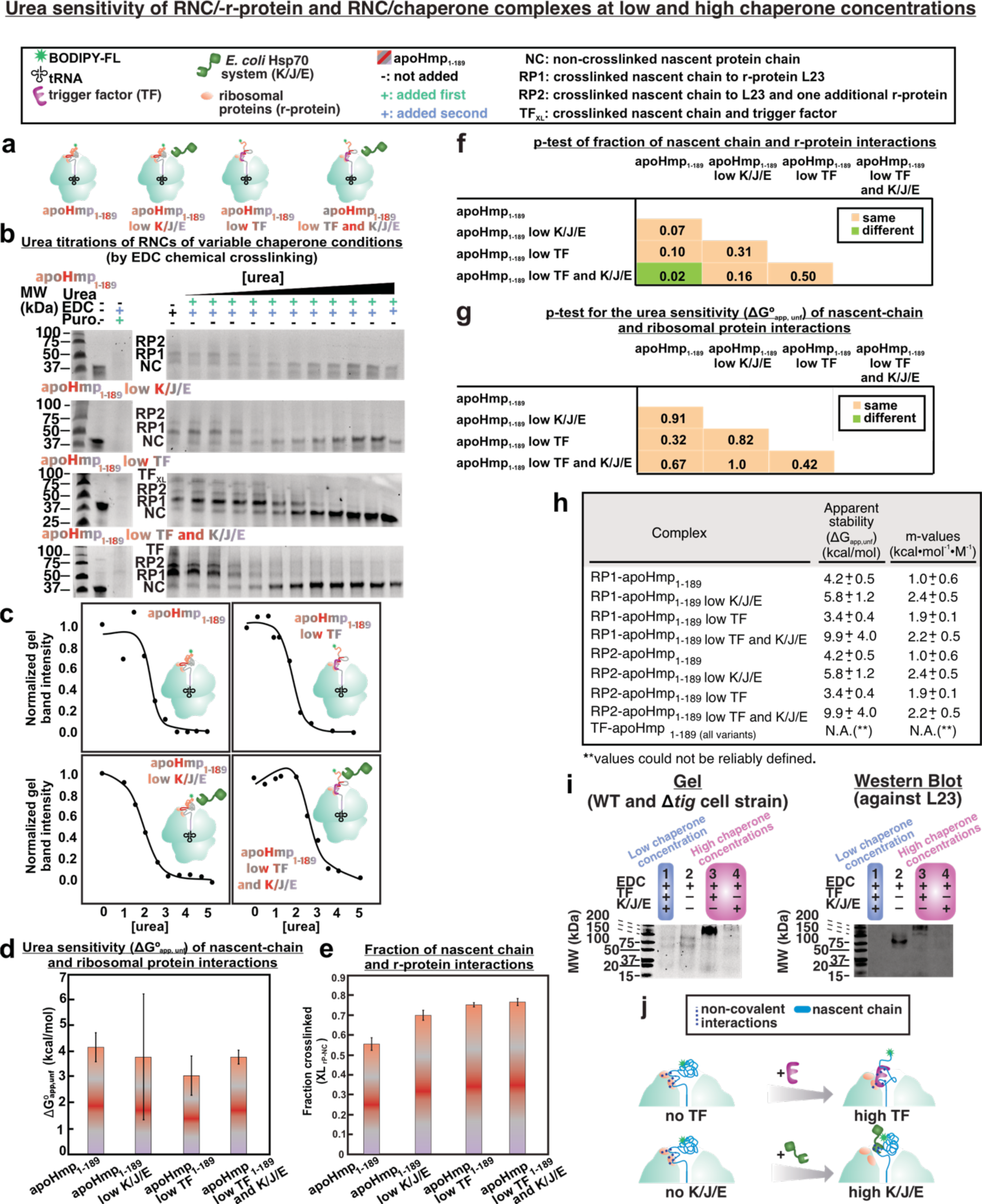
Low-pH SDS-PAGE and urea-titration analysis of ApoHmp_1-189_ in the absence and presence of TF and K/J/E chaperones. **a)** Cartoon illustrating the tested RNC constructs. **b)** Low-pH SDS-PAGE analysis showing apoHmp_1-189_ RNCs in complex with either r-proteins or molecular chaperones as a function of urea concentration. **c)** Representative urea titration curves. **d)** ΔG°_app,unfold_ values in the absence and presence of low (2-15 nM TF and 0.5 µM, 0.04µM and 0.05µM DnaK, DnaJ and GrpE respectively) concentrations of molecular chaperones. Error bars denote standard error based on 2-3 experiments. **e)** Fraction of RNC/r-protein complexes relative to total RNCs. Error bars denote standard error for 2-3 experiments. **f)** P-value table for two-tailed Student’s test assuming unequal variances (Welch’s t-test), comparing ΔG°_app,unfold_ values. Green and orange boxes denote statistically different and equivalent data, respectively, according to a ≥ 95% confidence interval. **g)** P*-* value table reporting on the fraction of RNC/r-protein complexes. Statistical assessments were similar to those listed in panel f. **h)** Table showing apparent unfolding free energies ΔG°_app,unfold_ and m values of relevant complexes. **i)** Low-pH SDS-PAGE analysis of apoHmp_1-189_ RNCs treated with EDC in the absence and presence of chaperones at low (lanes 1 and 2, (2-15 nM of TF and 0.5 µM, 0.04µM and 0.05µM of and DnaK, DnaJ and GrpE respectively) and higher concentrations (8 μM for TF and 10/2/4 μM for K/J/E respectively, lanes 3 and 4). Western Blot analysis probing interactions with ribosomal protein L23. **j)** Schematic representation of RNC interaction network at low (2-15 nM TF, and 0.5 µM, 0.04µM and 0.05µM DnaK, DnaJ and GrpE, respectively) and higher (8 μM and 10/2/4 μM K/J/E, respectively) chaperone concentrations.

These results show for the first time that two RNCs lacking signal or arrest sequences and encoding nascent proteins that partially fold on the ribosome experience interaction with a specific ribosomal protein. The intrinsically disordered PIR RNC, which was studied before (48), was found to have a ca. 50% population of RNC interacting with L23 and L29. In contrast, here we use two nascent chains corresponding to the foldable apoHmpHa_1-140_ and apoHmp_1-189_ sequences, whose N-terminal regions are compact (Fig. 1 c-g). These apoHmpH RNCs interact only with L23 and not with L29.

The interactions experienced by apoHmp_1-140_ and apoHmp_1-189_ involve the dominant fraction of the apoHmpH RNC population, which is found to nearly-quantitatively crosslink with the L23 ribosomal protein (Fig. 4 and 5). This scenario is different from the case of intrinsically disordered PIR RNCs, (48) which crosslink only in part, under the same experimental conditions. The larger extent of crosslinking of the foldable apoHmp_1-140_ and apoHmp_1-189_ nascent chains, however, may simply result from the greater number of EDC-reactive residues of apoHmp_1-140_ (25 EDC-reactive residues, 20 beyond the tunnel core) and apoHmp_1-189_ (36 EDC-reactive residues, 30 beyond the tunnel core) relative to PIR (14 EDC-reactive residues, 12 beyond the tunnel core). Interestingly, as shown in the sections below, the urea sensitivity of the L23-RNC complexes is similar for all RNCs, suggesting comparable interaction strengths.

Importantly, given that the fluorescence anisotropy-decay data of Figure 1 show that RNCs of both apoHmp_1-140_ and apoHmp_1-189_ have “freely tumbling” compact regions, our data suggest that the compact regions are not engaged in direct interactions with the ribosomal surface.

In the case of apoHmp_1-189_, there is an additional interacting population, which we denoted as RP2 (Fig. 5). This population includes r-protein L23 (Fig. 5b) and one additional unidentified protein of c.a. 6-10 kDa. Our Western Blots indicate that L29 (7 kDa) is not present in the RP2 band (SI Appendix, Fig. S8b), based on molecular-weight arguments. Other cytoplasmic *E. coli* chaperones and ribosome interactors (GroEL, GroES, SecB, K/J/E, SRP and ClpB; MW range: 48-80 kDa) are ruled out as they would appear well above the RP2-containing band in our gels (Fig. 5). Because of its close spatial proximity to L23 (Fig. 4a) and based on molecular weight arguments, we believe that it is possible that RP2 comprises L23 and L29. However, our monoclonal antibodies were not able to capture an L29 epitope, suggesting the utility of experiments employing polyclonal L29 antibodies. These are experiments are planned in future investigations.

Finally, the data for Figures 4 and 5 also show that a fraction of the RNCs interacts with the trigger factor (TF) chaperone (presumably coupled to the ribosome via L23), rather than with theL23 r-protein. The role of the interactions between apoHmpH RNCs and TF are beyond the scope of this work and have already been explored elsewhere in the case of other client proteins. (39, 81–84) These studies showed that TF interacts with clients that bear an expanded conformation in their bound state.

In all, our data show that apoHmpH_1-140_ and apoHmp_1-189_ RNCs interact with either the L23 r-protein or with the TF chaperone. We propose that these two classes of interactions may play a similar role, and that therefore L23 may be a chaperone-like ribosomal protein that contributes to maintaining RNC solubility during translation. Future studies will focus on genetic r-protein modifications aimed at disrupting the detected interactions. The large surface-exposed nonpolar patch of the L23 ribosomal protein (Fig. 4b) suggests that the interactions involving apoHmp_1-140_ and apoHmp_1-189_ RNCs may be contributed by the hydrophobic effect. This nonpolar patch appears an ideal candidate for genetic modifications aimed at significantly perturbing RNC/r-protein interactions.

### Nascent chain-L23 complexes have the same apparent stability regardless of RNC sequence

To further explore the nature of the interactions between the L23 r-protein and nascent chains of increasing length and variable sequence, we performed urea titrations with chemical crosslinking detection (Fig. 6a,b). EDC readily reacts with amines and carboxylic acid functional groups, and there is no loss of EDC reactivity even in the presence of high urea concentrations (85). It is worth noting that the interactions identified in this work are not induced by the covalently N-terminal-linked BODIPY-502 fluorophore, as previous work has shown that this fluorophore does not interact with resuspended ribosomes under conditions similar to those of the present study (17). Therefore, by unfolding the complex in the presence of urea and subsequently adding EDC, we expect to gain insights into the urea sensitivity of nascent chain-L23 complexes. After collecting gel data on representative apoHmp and PIR nascent chains (Fig. 6c), we estimated the apparent stability (ΔG°_app, unfold_) of nascent chain-L23 complexes following procedures similar to the other urea titrations described earlier (76). Representative EDC-mediated urea titrations are shown in Figure 6c. Corresponding plots and apparent-stability data are displayed in Figure 6e,f. The matching two-tailed Student’s t-test is provided in Figure 6g. As shown in Figure 6h, the apparent stability values for the apoHmp and PIR nascent-chain/L23 complexes range between ca. 2 and 5.5 kcal/ mol. Remarkably, all complexes display the same urea sensitivity within error. This is true even though the nascent-chain portions that are not buried within the exit-tunnel core have widely different nonpolar and net-charge-per-residue content (Fig. 6i) and widely different total nonpolar surface accessible surface-area values (Fig. 6j).

In summary, the urea-titrations in Figure 6 show that the urea sensitivity of r-protein-nascent-chain complexes is similar regardless of the nature and length of the nascent chain, across the short and long (55- to 189-residue) chains examined here. In other words, RNC/-r-protein complexes have the same apparent stability although the RNCs have widely different physical properties and compaction, as discussed above and, in the case of PIR lack of compaction, in previous work. (48, 49) A cartoon showing a low-resolution RNC model consistent with the crosslinking and fluorescence-anisotropy data is shown in Figure 1c-g. Further, the longer RNCs of apoHmp_1-189_ also exhibit interactions with an additional yet-unidentified r-protein. The corresponding crosslinked complex is denoted as RP2 in Figure 5. At this juncture, a few additional considerations come to mind. First, the 55- and 140-residue nascent chains of apoHmp lack or include the N-terminal 65-94-residue compact region detected via fluorescence anisotropy decay (Fig. 1c-g), respectively. Therefore, the non-compact C-terminal portion of the nascent chain out of the tunnel core seems to be primarily engaged in the detected interactions with the L23 r-protein in the case of apoHmp. Given that the amino-acid sequences of the interacting regions of apoHmp_1-55_ and apoHmp_1-140_ must be different yet the interactions are of comparable apparent strength, the interactions seem to be of nonspecific nature, amounting to an overall sufficient affinity the 100% interacting population detected in the gels of Figure 6. Second, apoHmp_1-140_ and apoHmp_1-189_ are also found to experience interaction of equivalent apparent strength with L23 (see RP1 gel bands in Fig. 6d and g). On the other hand, apoHmp_1-189_ also experiences interactions with another r-protein (see RP2 band in Fig. 6d). This finding supports the idea that the RNC-r-protein contacts are of somewhat non-specific nature and longer RNCs interact with additional ribosomal surface, likely engaging a longer portion of their chain. Finally, previous work showed that the interactions experienced by PIR_1-91_ are ca. 50% mediated by Mg^+2^ ions (48). Yet the apparent strength of these interactions is not different from that experienced by the other nascent chains, suggesting that all interactions are relative weak and non-specific (Fig. 6 f-h). This model is consistent with the fact RNC-r-protein interactions likely need to be continuously remodeled, as nascent chains elongate during translation. In all, the nature (Mg^+2^ dependence, nonpolar *vs* electrostatic, or else) of RNC-r-protein interactions is still poorly understood and clearly awaits future higher-resolution structural investigations.

### Nascent chain and r-protein interaction strength does not vary in the presence of one or more molecular chaperone

Next, we explored the effect of molecular chaperones TF and Hsp70 on the RNC-r-protein interactions via the same type of EDC-mediated urea titrations employed in the last two sections. The Hsp70 chaperone was examined holistically as the as Dnak/DnaJ/GrpE chaperone system, denoted here as K/J/E. TF is known to associate with prokaryotic ribosomes (86) and K/J/E works in cooperation with TF (87) to promote nascent-protein folding and prevent their aggregation (88–91).

First, we evaluated apoHmp_1-189_ devoid of both TF and the Hsp70 chaperone system (K/J/E), apoHmp_1-189_ in the presence of low concentrations of TF only, apoHmp_1-189_ in the presence of low concentrations of K/J/E only, and apoHmp_1-189_ in the presence of both chaperones at low concentration (Fig. 7a). Urea titrations were carried out using increasing concentrations of urea (Fig. 7b), and the intensity of the crosslinked fraction was plotted as a function of urea (Fig. 7c). We then obtained a ΔG°_app, unfold_ values for each of these constructs (Fig. 7d) and evaluated them with a two-tailed Student’s t-test (Fig. 7g). Interestingly, in all construct variations tested, the apparent strength of the L23-nascent chain complex was found to be statistically similar (Fig.7f, h). Because this effect is not due to an increase or decrease in the fraction of crosslinked nascent chains to r-proteins (Fig. 7 e, f). Via Western Blot analysis, we were able to conclude that interactions mainly take place with r-protein L23 (Fig. 5, b and SI Appendix, Fig. S8). This finding is consistent with our assessment of interacting proteins from apoHmp_1-140_ (Fig. 4 and SI Appendix, Fig. S5). This result suggests that in both the presence and absence of one or more molecular chaperones, nascent chains interact with ribosomal L23 in a structurally similar manner, though future work will need to be employed to validate this claim.

To further elucidate the nature of nascent chain and molecular chaperones, we performed experiments using increasing concentrations of TF and K/J/E in the presence of EDC (Fig. 7i, j). Briefly, interactions with r-protein L23 can be displaced by high concentrations of molecular chaperones (8 μM TF and 10/2/4 μM K/J/E, respectively, Fig. 8c,d). A low pH SDS-PAGE gel reporting on apoHmp_1-189_ RNCs shows that at low concentrations of TF and K/J/E (2-15 nM of TF and 0.5 µM, 0.04µM and 0.05µM of and DnaK, DnaJ and GrpE respectively), interactions with two r-proteins are established (Fig. 7i, gel lanes 1 and 2). Corresponding lanes in the Western Blot of this gel against L23 show that interactions involve r-protein L23 (Fig. 7i, Western Blot lanes 1 and 2). Interestingly, at high TF concentration (8 μM) Fig. 7i, gel lane 3), all r-protein interactions are displaced and replaced by contacts with TF (Fig. 7i, Western Blot lane 3). TF and L23 are known to interact with one another on the ribosome,(81, 83, 91) though we presently cannot explicitly discriminate whether the nascent chains interacts with L23 and TF, or if the nascent chain interacts with TF, which in turn interacts with L23. Here, we propose the simplest scenario namely that RNCs interact with TF only and, in turn, TF interacts with L23, which is known to be the TF docking site on the ribosome (2).

**Figure 8.**
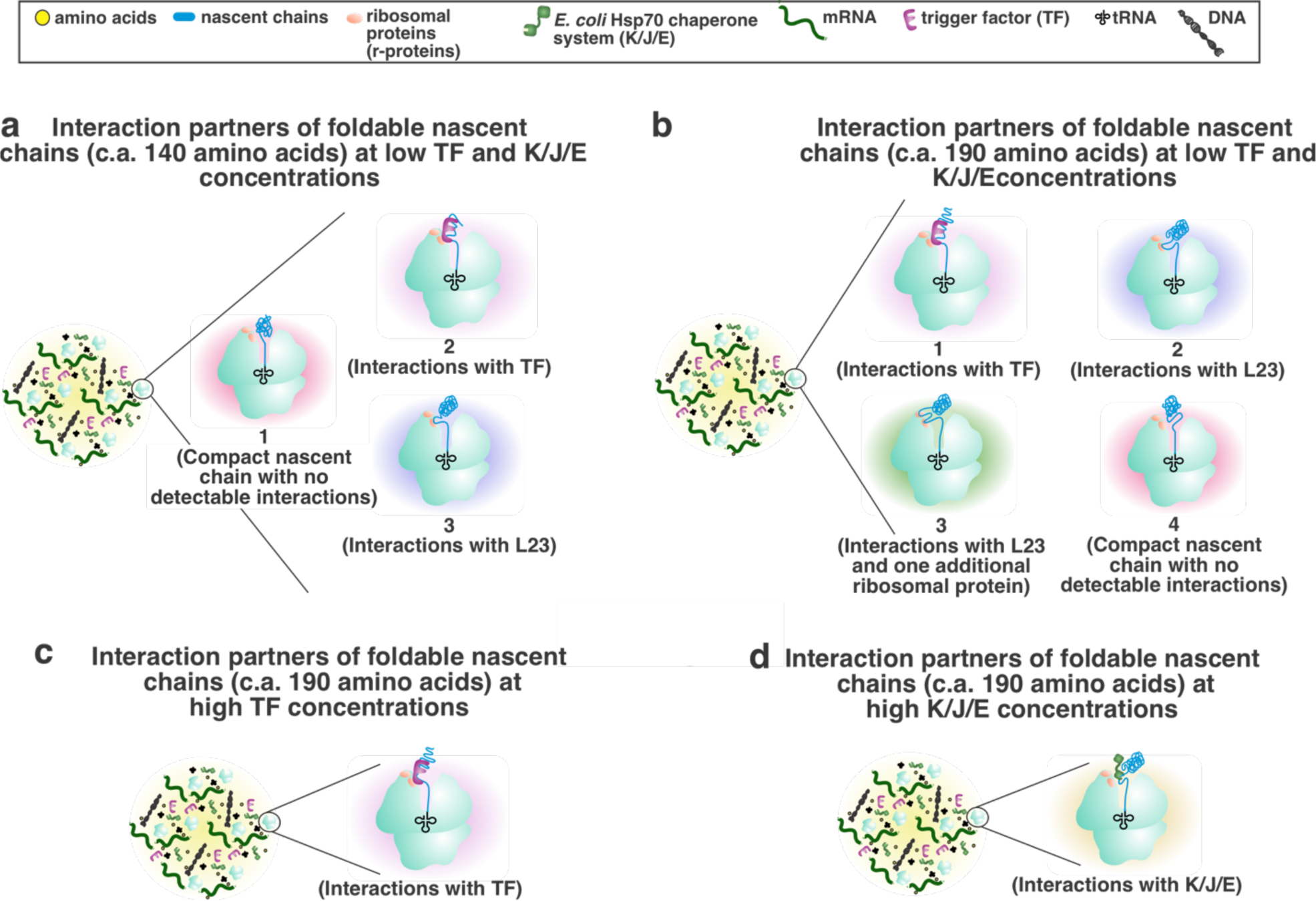
Cartoon summarizing the RNC interaction networks detected in this work under different experimental conditions. Interaction partners of **a)** apoHmp_1-140_ and **b)** apoHmp_1-189_ RNCs at low chaperone concentrations (2-15 nM of TF and 0.5 µM, 0.04µM and 0.05µM of and DnaK, DnaJ and GrpE respectively). **c)** Interaction partners of apoHmp_1-189_ RNCs at higher **c)** TF (8 μM) and **d)** K/J/E chaperone concentrations (10/2/4 μM K/J/E, respectively).

Similarly, interactions with r-proteins are also displaced by increased concentrations of K/J/E (10/2/4 μM K/J/E, respectively), although no interactions with L23 (Fig. 7i Western Blot lane 4), L24 or L29 (SI Appendix, Fig. S9) were detected when K/J/E concentrations are increased, suggesting that the nascent chain likely does not interact with any r-proteins (Fig. 7i Western Blot lane 4). Similar results are seen for apoHmp_1-140_ (SI Appendix, Fig. S7). Figure 8b and c shows model of the interaction interplay between nascent chains (c.a. length of 140-190 amino acids), r-proteins and molecular chaperones assessed in this work.

To summarize, when TF and K/J/E are not present, the nascent chain interacts with r-proteins only. In experiments with low concentrations of molecular chaperones (2-15 nM of TF and 0.5 µM, 0.04µM and 0.05µM of and DnaK, DnaJ and GrpE respectively, Fig. 7i,j), RNCs interact with TF but not K/J/E. At high, physiologically relevant concentrations of TF and K/J/E (8 μM TF and 10/2/4 μM K/J/E, respectively), RNC interactions with r-proteins are displaced by interactions with molecular chaperones. It is worth nothing that TF and K/J/E chaperones are shared with thousands of additional cellular proteins *in vivo,* unlike in the experiments shown here, which include purified resuspended RNCs. Further, our RNC concentrations are only 20-30 nM. Hence, chaperones are in large excess over RNCs even at the low chaperone concentrations employed here. This scenario differs from the cellular environment where both RNCs and molecular chaperones are at comparable concentrations, within the low uM range. Therefore, we propose that the actual cellular milieu likely involves RNC populations that interact in part with TF and K/J/E and in part with r-proteins.

### Concluding remarks on the RNC interaction network experienced by the foldable apoHmpH protein

An overall model recapitulating the main features of the RNC interactions network elucidated in this work in shown in Figure 8. Briefly, RNCs encompassing the entire N terminal domain of apoHmpH (apoHmp_1-140_) experience either interactions with the L23 r-protein accompanied by independent tumbling of the N-terminal region or interactions with the TF chaperone (Fig. 8a). On the other hand, RNCs comprising the entire N terminal domain of apoHmpH and additional 49 residues belonging to the second domain (apoHmp_1-189_) are subject to either similar interactions with the L23 r-protein, or to interactions with L23 and one additional presently unidentified protein (RP2), or to interactions with TF (Fig. 8b). Finally, in the presence of an excess of either TF or the Hsp70 chaperone system at total concentrations matching cellular levels, the interactions with ribosomal proteins are replaced by interactions with the respective molecular chaperones (Fig. 8c-d). The emerging scenario resulting from our work suggests that the ribosome plays an RNC-interacting role entirely similar to that of molecular chaperones.

In all, our findings highlight the prominent role of the ribosome as an RNC interactor, and suggest that the ribosome may have played a primordial chaperone role in Nature, before the evolution of molecular chaperones.

## Materials and Methods

### Preparation of empty ribosomes

Empty ribosomes were generated from an in-house**-**prepared A19 WT or A19 Δtig *E. coli* S30 cell extract as described (17, 92). Briefly, cells were grown in Luria-Bertani (LB) broth and harvested at mid-log phase (A_600_ ∼ 0.6). The cells were lysed through a French press (thermo Electron Corporation, Waltham, MA) at ∼12,000 psi with a single passage. The lysate was subject to centrifugation at 30910 g and 20 °C for 20 min. After centrifugation, the supernatant was incubated in translation buffer (0.75 M Tris-HCl pH 8.2, 7.5 mM DTT, 21 mM Mg(OAc)_2_, 500 µM amino acids, 6 mM ATP, 67 mM PEP and 160 µg/mL pyruvate kinase) for 80 min to remove any endogenous mRNA from ribosomes. The supernatant was then dialyzed (12-14 kDa MWCO) in buffer (10 mM Tris-HCl pH 8.2, 14 mM Mg(OAc)_2_, 60 mM KOAc and 1 mM DTT) for 12 hrs, with a buffer exchange every 4 hours. The resulting A19 cell extract was used as the empty-ribosome sample.

### Preparation of RNCs

RNCs were generated using an in-house prepared A19 *E. coli* transcription-translation coupled cell-free system (17, 92) as described. Cell strains either including (WT) or lacking (*Δtig*) the trigger factor gene were employed (17, 92). Hsp70 chaperone activity was suppressed via the KLR-70 peptide (93) to a final concentration of 0.2 mM. Transcription-translation proceeded for 30 min at 37 °C in the presence of BODIPY-FL-Met-tRNA^f-Met^ to specifically label RNCs at the N terminus. BODIPY-FL-Met-tRNA^f-Met^ was prepared as described (17). RNCs were stalled at various lengths to generate the desired apoHmp and PIR constructs via oligodeoxynucleotide-directed mRNA cleavage (17, 94, 95). An anti-ssrA oligonucleotide (17) was added to a final concentration of 12.83 pmol/µL to prevent premature release of stalled RNCs. RNC pellets were isolated via a sucrose cushion (1.1 M sucrose, 20 mM tris base, 10 mM Mg(OAc)_2_, 500 mM NH_4_Cl, and 0.5 mM EDTA, 1 mM DTT, pH 7.0, as described)(17) and subjected to ultracentrifugation at 160,000 g for 1 hr at 4 °C. The purified pellet was dissolved in resuspension buffer (10 mM tris-HCl, 10 mM Mg(OAc)_2_, 60 mM NH_4_Cl, 0,5 mM EDTA and 1.0 mM DTT, pH 7.0) by shaking in an orbital shaker at 200 rpm on ice for 1 hr.

### Other experimental procedures

Details on sucrose gradients, low-pH gels, puromycin assays, chemical crosslinking and urea titrations are available in the SI Appendix.

## Acknowledgments

We are thankful to M. Dalphin for helpful discussions. This work was funded by the National Science Foundation (NSF) grants MCB-1616459 and MCB-0951209 (to S.C). M. M. M. and R.B.H. received NIH TEAM-Science Fellowships from the University of Wisconsin-Madison and M.M.M received the Straka Fellowship from the University of Wisconsin-Madison. A. E. V. received a National Science Foundation GRFP graduate fellowship and a Science and Medicine Graduate Research Scholars Fellowship from the University of Wisconsin-Madison.

## SI APPENDIX

### SUPPORTING TEXT

#### Sucrose-gradient analysis of 70S empty ribosomes and ribosome-peptidyl-tRNA complexes

In order to explore the effect of nascent-chain characteristics on the bacterial ribosome, we examined whether the incorporation of aminoacyl initiator tRNA (Met-tRNA^fMet^) or very short peptidyl tRNAs affects the ribosomal complex in the presence of urea. We collectively denote these species, which were generated via oligodeoxynucleotide-directed mRNA cleavage (17, 94, 95) (see Methods), as tRNAs carrying short nascent chains, or snc-tRNAs. Note that the antisense DNA construct used in the oligonucleotide-directed mRNA cleavage approach was designed to generate ribosome stalling after the first N-terminal residue (Met) of the nascent chain only. It is known that the *E. coli* RNAse H enzyme, employed here in conjunction with antisense oligodeoxynucleotides, typically establishes well-defined sharp cleavage sites, characterized by a site-specific distribution of the cleavage site of ca. 1-3 nucleotides (95). Hence a dominant population of very short (1-3 residues) nascent chains in the ribosomal complexes encompassing snc-tRNAs is expected, consistent with the observed sharp gel bands (SI Appendix, Fig. S1). It follows that the nascent chains belonging to snc-tRNAs are not sufficiently long to interact with ribosomal proteins (r-proteins) across the ribosomal exit tunnel (1). In addition, the specific thermodynamic stabilization imparted by RNA-oligodeoxynycleotide complexation was designed to be at least 12.4 kcal/mole, with a corresponding antisense-oligodeoxynucleotide length ranging from 11 to 38 DNA bases. The above free-energy value is significantly larger than what was used in the original report of the oligodeoxynucleotide-directed RNA cleavage approach (95). Therefore, ribosomal complexes that include snc-tRNAs are expected to bear no contributions due to interactions of nascent chains with ribosomal proteins. It is worth noting, however, that in the case of some specific RNA sequences under non-optimal conditions, the cleavage-site distribution width was found to be larger than 1-2 residues (98). Therefore, in our case, any hypothetical longer RNCs than 1-3 residues are expected to be poorly populated.

Once appropriate RNCs harboring snc-tRNAs were made, we set out to test the response of the ribosome against exposure to urea. Specifically, we compared the urea sensitivity of empty 70S ribosomes (Fig. 2a-b) to that of ribosomes harboring snc-tRNAs (Fig. 2c) by sucrose gradient-detected urea titrations. Sucrose gradients are able to resolve ribosomal subunits and entire 70S ribosomal particles.(99) Further, these gradients have previously been employed to monitor the unfolding of whole ribosomes or ribosomal subunits in the absence and presence of targeted structure-perturbing buffers (57, 60, 100–105) (e.g., containing EDTA or other related agents) or classical denaturing agents like urea.(33, 64, 106) Conveniently, high concentrations of urea do not perturb elution-profile integrity through line-broadening or other effects. (107) Therefore, urea titrations of the 70S ribosome including sucrose-gradient detection are a powerful tool to explore ribosomal-subunit dissociation and unfolding.

Sucrose-gradient elution profiles were monitored at 260 (Fig. 2) and 280 nm (SI Appendix, Fig. S3 and S4) in separate experiments, to probe for any potential differences in the response of rRNA and r-proteins. Our data show that rRNA and r-proteins belonging to 70S ribosomal particles devoid of tRNA and nascent chains, denoted here as empty ribosomes, are overall stable up to 1.0 M urea, with only a small extent of dissociation of the 70S particle into its 30S and 50S subunits (Fig. 2a). At 2 M urea, empty ribosomes undergo subunit dissociation accompanied by extensive line-broadening. Due to the dominant presence of intact 16S and 23S rRNA band in ethidium bromide-stained agarose gels (Supporting Fig. S2), line broadening is not ascribed to rRNA degradation. Therefore, we interpret the broad sucrose-gradient peaks observed at 2 M urea as diagnostic of rRNA and(or) r-protein unfolding accompanied by conformational heterogeneity. At > 2 M urea, both the empty ribosome, and ribosomes harboring snc-tRNA undergo severe line-broadening (Fig. 2). At the highest urea concentration tested in this work (4 M), line-broadening is so extensive that it is difficult to deconvolute the contribution of individual ribosomal components. A similar scenario is supported by the empty-ribosome data at 280 nm, suggesting that rRNA and r-protein unfolding proceeds in concert (Supporting Fig. S3).

Ribosomes carrying tRNAs linked to very short nascent chains (snc-tRNA) are also characterized by progressive disassembly, as urea concentration increases (Fig. 2 and Supporting Fig. S4). Unlike empty ribosomes, however, the ribosomes harboring snc-tRNAs are still ca. 50% intact at 2 M urea. Further, at this urea concentration the 30S and 50S subunits (dissociated from 70S intact ribosomes) have not yet undergone any unfolding, given that no line broadening is observed.

Notably, the sucrose gradient profiles of ribosomes harboring snc-tRNAs also display two additional peaks eluting after the 70S ribosome (Fig. 2). These features are either due to polysomes (i.e., nearby ribosomes linked via the same mRNA strand) or to self-associated ribosomes brought together by through-space noncovalent surface contacts. The mRNA encoding short nascent-chains (sncs) is predominantly cleaved only 11 ribonucleotides away from the mRNA ribosome binding site, i.e. the Shine-Dalgarno sequence (108). Ribosome profiling (109) and structural considerations reveal that each translating bacterial ribosome spans a length corresponding ca. 24 ribonucleotides (110). Further, computer simulations suggested that polysomes have c.a. 24 residues between neighboring ribosomes (111). Therefore, geometrical factors render polysome formation very unlikely. To gain further insights, transmission electron microscope (TEM) negative stain images of ribosomes carrying snc-tRNAs were acquired in the presence of 2% methyl-tungstate (112). The representative TEM image displayed in Figure 3 shows that, in addition to isolated ribosomes (within dashed blue squares), some closely spaced ribosomes (within dashed red squares) comprising 2 or more particles are also present. The spatial arrangement of these particles renders it impossible to establish where these species are polysomes or other forms of self-associated ribosomes. While polysomes seem unlikely due to the above-listed geometrical arguments, it is possible that some longer-than expected nascent chains may be populated in these samples, preventing the ruling out of polysomes. In all, the origin of the late-eluting peaks found in ribosome samples harboring snc-tRNAs remains unestablished and further future investigations are required to shed further light on this matter. On the other hand, the disassembly pattern of these peaks is entirely similar to that of 70S ribosomes harboring snc-tRNAs. Therefore, the late-eluting peaks do not add any new information nor modify the conclusions reached for the 70S particles.

The urea sensitivity of a representative ribosome harboring a longer nascent chain (32- residue long) derived from *E. coli* Hmp_108-140_ has also been analyzed. The results are shown in Figure 4 and SI-Appendix Figure S3. These ribosomes behave in the same way as ribosomes harboring snc-tRNAs, suggesting that nascent-chain length does not affect the apparent stability of the 70S ribosome, and that the enhanced apparent stability may be dominated by the role of the tRNA. Additional experiments probing this topic in further detail are described in some sections of the main manuscript.

In summary, the data in Figures 2-4 show that empty-intact 70S ribosomes are more sensitive to urea-induced denaturation than ribosomes carrying tRNAs linked to nascent chains of variable lengths.

### SUPPORTING MATERIALS AND METHODS

#### Denaturation of ribosome-nascent-chain complexes and empty ribosomes

A 10 M stock solution of 0.22 µm-filtered urea was prepared in resuspension buffer and the refractive index was measured with an Abbe Refractometer (Thermo Spectronic, Fisher Scientific) to derive actual urea concentrations. RNCs subject to sucrose cushion ultracentrifugation (see section on RNC preparation) or empty ribosomes obtained from crude S30 were incubated in the presence of variable concentrations of urea for 1 hr at ambient temperature in the dark.

#### Assessment of urea sensitivity of 70S ribosome and RNCs via sucrose gradient analysis

An in-house prepared A19 *E. coli* mixture (with 70S ribosomes) was incubated for 1 hr at variable urea concentrations at ambient temperature. Samples were loaded onto a 5-45% sucrose gradient and centrifuged using a Beckman L-70 Ultracentrifuge with a SW41 rotor at 288, 000 x g for 1.5 hrs at 4 °C. Gradient profiles were obtained on a Biocomp Fractionator at 0.2 mm/sec. The absorbance was measured at 254 nm and 280 nm using a Triax flow cell from BioComp to check for intactness of the rRNA and r-proteins, respectively. RNCs were treated in a similar manner after denaturation in urea for 1 hr (see RNC denaturation in Methods). The 30S, 50S and 70S subunits were collected in both instances and loaded on a 2% agarose gel (0.02 M Tris Base, 0.01 M acetic acid and 0.0005 M EDTA, or 0.5x TAE). Ethidium bromide was added to a final concentration of 0.5 µg/mL. Samples were run for 100 min at 3.92 V/cm. The gel was imaged on a GE FLA 9500 Laser Imager at PMT values between 500-700.

#### Imaging of RNCs via negative staining

RNC samples prepared in the presence of Met-tRNA^fMet^ (2 μL) were placed onto a glow-discharged copper 300-mesh formvar-carbon grid (made in house by Medical Sciences Electron Microscopy staff at UW-Madison), blotted with filter paper and allowed to dry. A Nano-W staining solution (Nanoprobes) was placed on the grid in equal volume, blotted with filter paper and allowed to dry. Images were collected on a CM120 transmission electron microscope (Philips) at 140000x magnification and 80 keV using a BIOSPR12 camera.

#### Assessment of apparent stability of RNC/-r-protein and RNC/chaperone complexes via a crosslinking assay

The EDC crosslinker is capable of capturing interactions involving RNCs and r-proteins or chaperones within the ribosomal exit-tunnel vestibule and outside the ribosomal exit-tunnel core (48). Either ribosome-bound or ribosome-released control samples (with EDC added after RNC ribosome release) were used to probe RNC production and to test the lack of ribosome-released nascent-chain crosslinking to other species. After incubation in urea for 1 hr, a 10x concentration of EDC solution (800 mM EDC, pH 6.8-7.0) was added to RNCs to a final concentration of 1x. Samples were incubated for 30 min at 30 °C and then quenched with a 10x concentration of Quenching Buffer (1.0 M Tris-HCl pH 7.0, 1.0 M Glycine, 1.0 M KOAc) to a final concentration of 1x (0.1 M Tris-HCl pH 7.0, 0.1 M Glycine, 0.1M KOAc). Samples were loaded onto a low-pH SDS-PAGE gels using either a 10% acrylamide gel (apoHmp_1-55_, apoHmp_1-140_ and apoHmp_1-189_) or a 9% acrylamide gel (PIR) in a 1:1 ratio with loading buffer. Gels ran at 3.92 V/cm for either 4 hours (apoHmp_1-55_, apoHmp_1-140_ and apoHmp_1-189_) or 2.5 hrs (PIR) and were imaged on a GE FLA 9500 Laser Imager. Fluorophores were excited at 473 nm, with a PMT value within the 700-950 range.

Crosslinked band intensities were evaluated via the ImageJ software (113). The normalized intensity of each band was assessed via relation 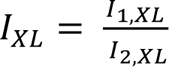, where I_XL_ is the normalized intensity of an individual crosslinked species, I_1, XL_ is the intensity of the individual crosslinked species, I_2, XL_ is the intensity of the species at 0 M urea. The fractional band intensities were then plotted as a function of urea concentration and then plotted to fit pre- and post-transition region baselines. A 2-state unfolding expression taking pre- and post-transition baseline slopes into account (76) was used to fit the raw urea-titration data and deduce m-value and ΔG°_H2O_ values via the Kaleidagraph software (77).

#### Western Blot analysis of r-protein interactions with L23, L24 and L29

Western Blots were performed as described (48). Aliquots of the rabbit anti-uL23 antibody were kindly donated by Shu-ou Shan (California Institute of Technology). The anti-uL23 antibody was generated by GenScript, using the CGKVKRHGQRIGRRS peptide as epitope and has been validated in previous work (114). Rabbit anti-uL17, -uL18/L22, -uL24, -uL29, and -uL32 antibodies were kindly facilitated by Bryan W. Davies (University of Texas-Austin) and Melanie Oakes (University of California, Irvine) who obtained them from Masayasu Nomura (University of Wisconsin-Madison). The antibodies were generated using the purified *E. coli* ribosomal proteins L17, L23, L18+L22, L24, L29, and L32 (115).

#### Assessment of apparent stability of PTC via a puromycin-assisted nascent-chain release assay

To confirm that the polypeptide was attached to the ribosome, a low-pH SDS-PAGE(75) using a 9% acrylamide gel was conducted before (positive control) and after treatment of 1 mM puromycin, which reacted with the samples for 30 minutes at 37 °C. Samples were loaded on gels in a 1:1 ratio with loading buffer. Identical samples of the positive control and puromycin-released samples at 2.23 M urea were loaded on each gel to control for intrinsic gel differences (i.e., gel crosslinking, which may affect the fluorophore quantum yield). Prior to gel loading, samples were heated at 37 °C for five min and allowed to sit at room temperature for 5 min. Gels ran at 3.92 V/cm for 3.5 hr and imaged on a GE FLA 9500 Laser Imager. Fluorophores were excited at 473 nm and a PMT value between 315-700.

Due to the number of samples, 2 gels were needed for the positive control samples and 2 gels were needed for the puromycin-released samples per experiment. Fluorescence intensity adjustments were made between gels by normalizing the 2.23 M urea samples from each gel via Equation 1

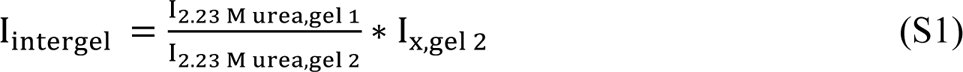

where I_2.23 M urea, gel1_ and I_2.23 M urea, gel2_ refer to the band intensities of the 2.23 M urea sample in each gel. I_gel2_ is the band intensity of the sample loaded onto the second gel and I_x,intergel_ is any given band intensity on the second gel in comparison to the band intensity of the first gel.

To control for differences in RNC concentrations and compare band intensities between the bound and released sample gels, Equation 2 was used

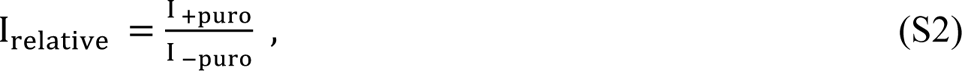

where I _+puro_ is the band intensity of the puromycin-released sample and I _-puro_ is the band intensity of the positive control. Band intensities were then divided by the band intensity of the sample containing the highest urea concentration to normalize intensities between 0 and 1. The resulting intensities were plotted to fit pre- and post-transition region baselines. Using an extrapolation method (76), the transition region slope and y-intercept were deduced to obtain the m-value and ΔG°_H2O_ with the Kaleidagraph software (77).

#### Ribosomal-protein (r-protein) stability assessment via a Trp fluorescence-emission assay

The *E. coli* r-proteins collectively contain 32 Trp residues. Trp is sensitive to changes in its environment and exposure to polar solvents causes a red-shift in its excitation spectra, which was monitored as a function of increasing urea concentrations (116–118). RNC samples were excited at 285 nm (bandpass of 4 nm) and the fluorescence was monitored from 295-500 nm (bandpass of 4 nm) on a Photon Counting Spectrofluorimeter (ISS) since the indole group of tryptophan is the major component of UV absorbance in that region (119).

To generate a titration curve for the RNC complex, the buffer spectra was first subtracted from the produced emission spectra. Then a baseline correction was done on the resulting spectra. The spectral center of mass of the resulting spectra was obtained using the emission spectra between 300-385 nm (to omit the scatter peak and Raman peak) and Equation 3

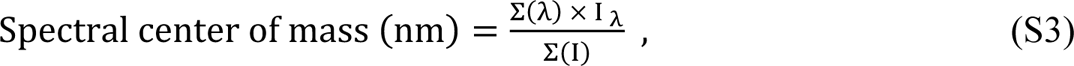

where λ is the wavelength and I_λ_ is the intensity at a specific wavelength.

The fraction of unfolded protein at each concentration of urea was determined according to

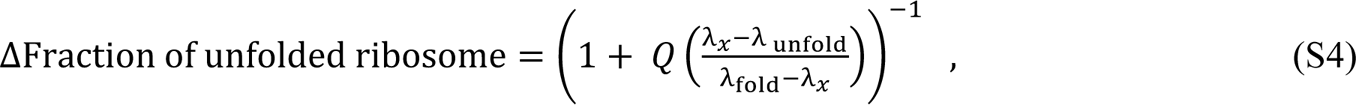

where λ_fold_ is the shortest wavelength calculated from Equation 3 for the folded species and λ_unfold_ is the longest wavelength calculated from Equation 3 for the unfolded species. Q denotes the ratio between the quantum yields of folded and unfolded states. Quantum-yield changes were calculated by taking highest intensity values from each folded sample (0.0 M, 0.13 M and 0.45 M urea) as well as each unfolded sample (5.5 M, 6.0 M and 6.5 M urea). The three folded and unfolded values were averaged amongst their respective groups to determine Q. ΔCoM was plotted as a function of urea concentration and the pre- and post-transition baselines were determined in Microsoft Excel. Free energy of unfolding curves were generated with the software Kaleidagraph (Synergy Software) using a known extrapolation method.

## SUPPORTING TABLES

**Table S1.**
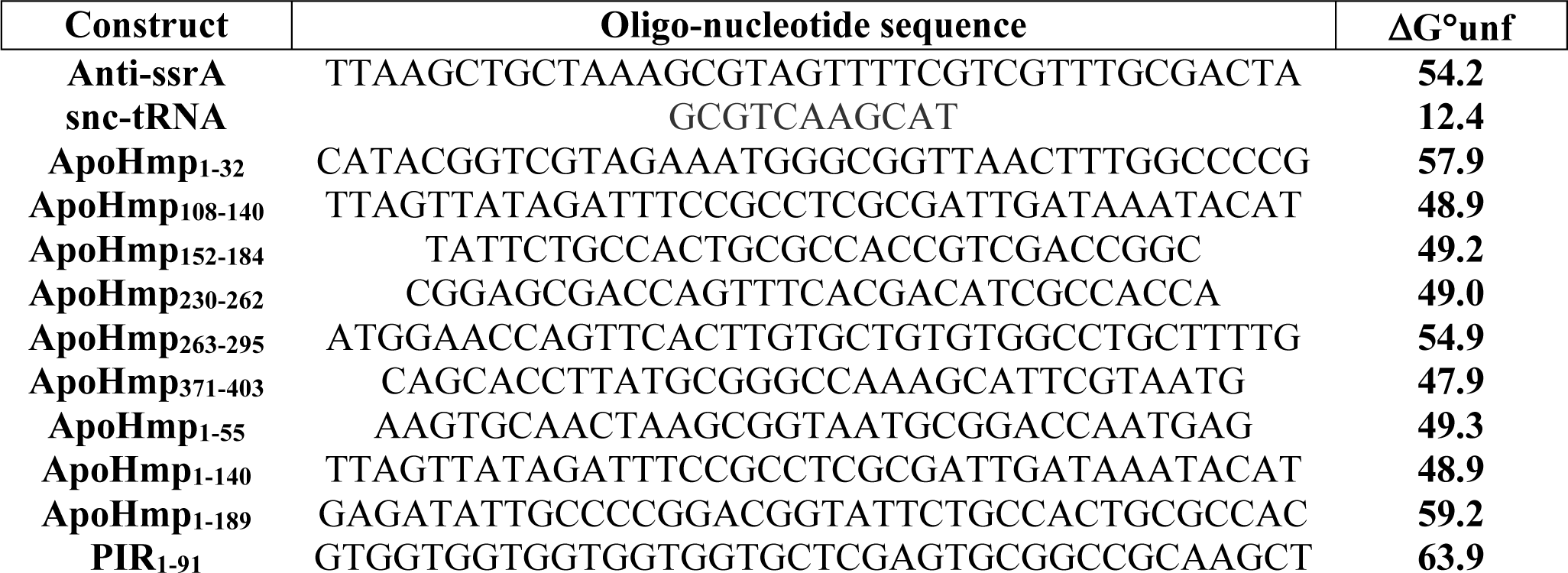
Name, nucleotide sequence, and calculated ΔG° of unfolding for each oligo nucleotide used in this work. ΔG° was calculated using an online calculator (biosyn.com). Sequences used for calculation are as shown.

## SUPPORTING FIGURE LEGENDS

**Figure S1.**
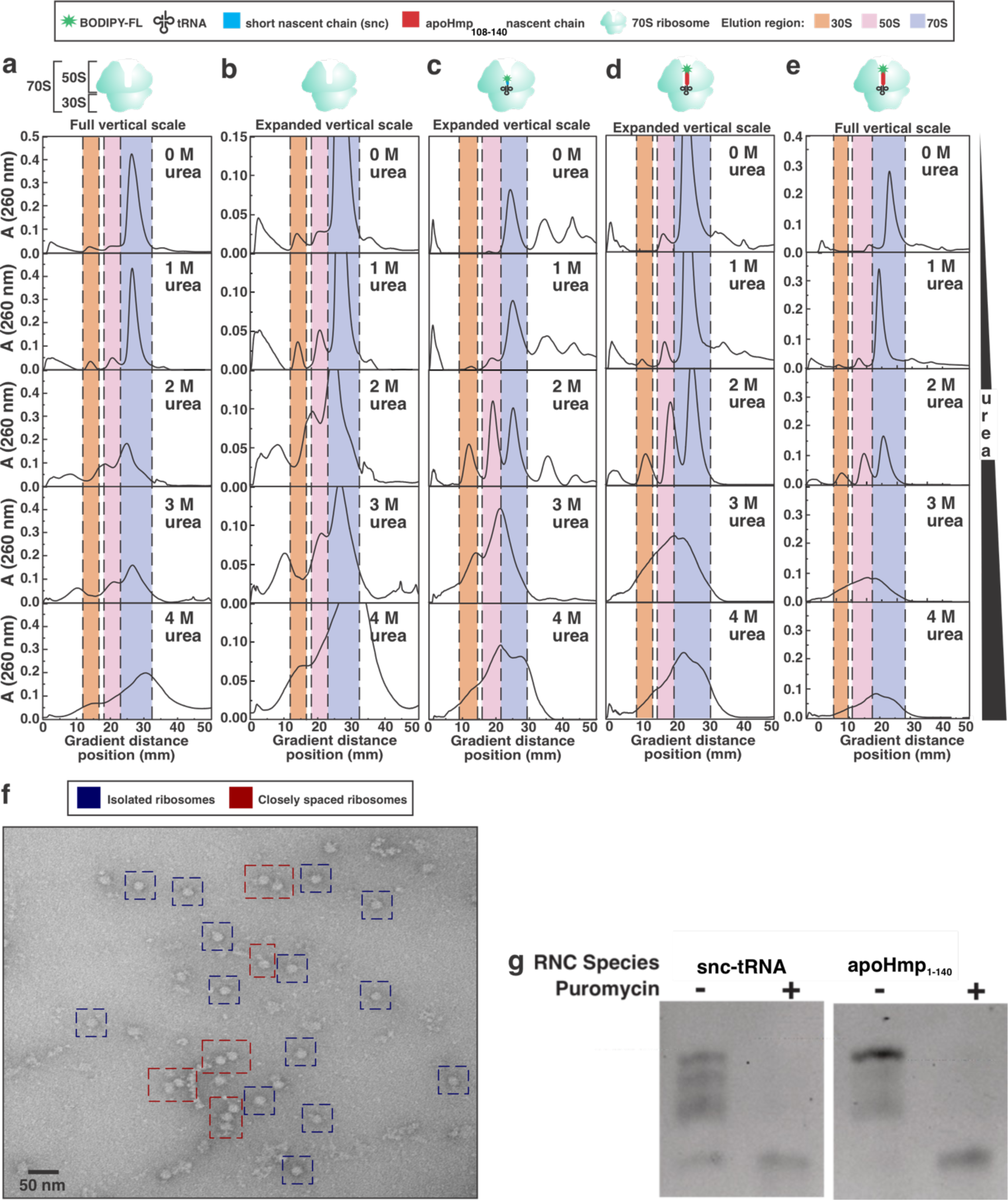
Sucrose gradient profiles for both empty 70S ribosomes (a), the same 70S ribosome profile is set to the same y-axis scale as the short (1-2 amino acid) nascent chains (snc-tRNA), (b) ribosomes harboring short (1-2 amino acid) nascent chains (c), ribosomes bearing a longer nascent-peptide chain (d), and the same ribosomes bearing a longer nascent-peptide chain at the full vertical scale (e) are shown for increasing concentrations of urea. Profiles were collected at 260 nm. (f) A TEM image of the snc-tRNA is shown. The dark blue squares show what are likely isolated 70S ribosomes and the red squares show closely spaced ribosomes. (g) Bound and released snc-tRNA and apoHmp_1-140_ nascent chains with the addition of puromycin (released) or without (bound) on a low pH 10% SDS-PAGE gel.

**Figure S2.**
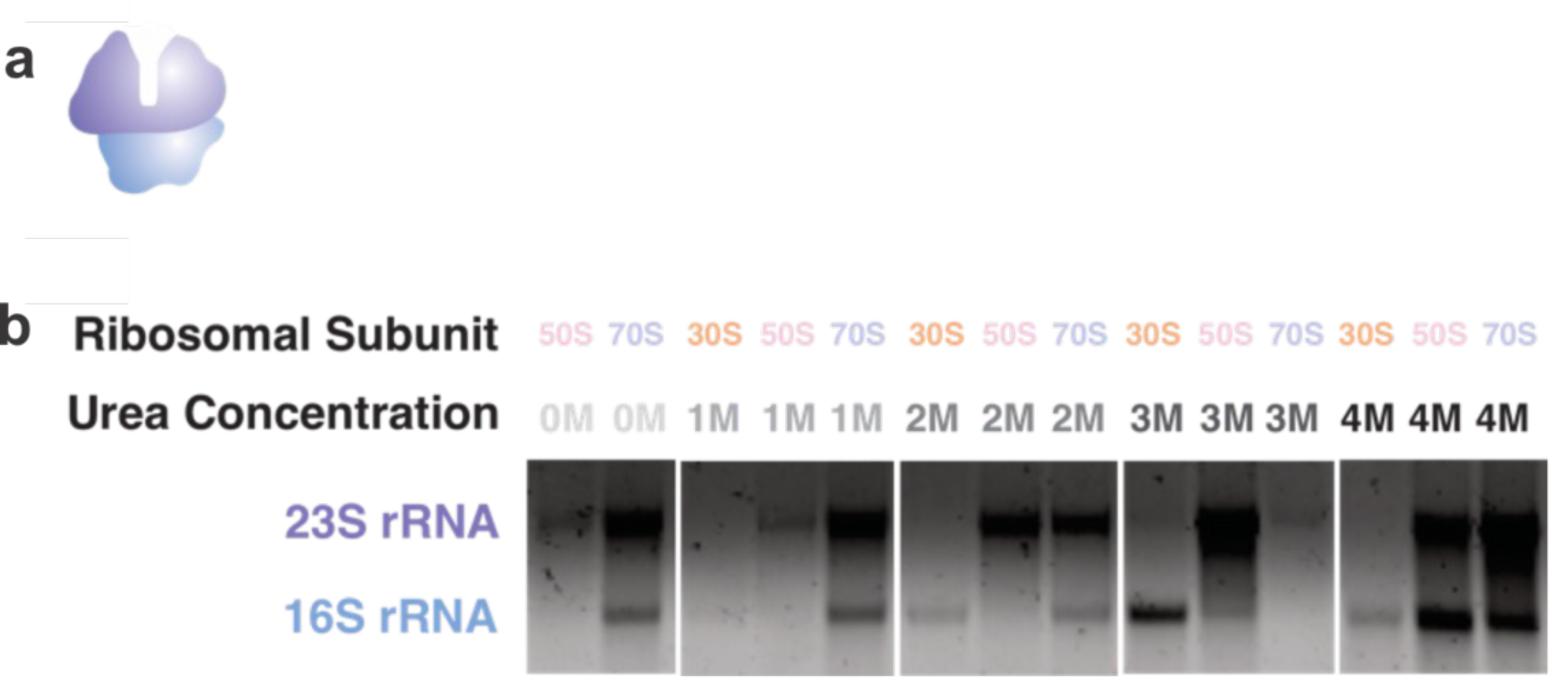
(a) The 70S bacterial ribosome contains two subunits, the 30S (blue) and the 50S (purple), which contain the 16S and 23S rRNA respectively. (b) Representative agarose gel stained with ethidium bromide for the 30S, 50S and 70S of ribosome samples at increasing concentrations of urea. Subunit peak corresponds to the area in which the sample was collected.

**Figure S3.**
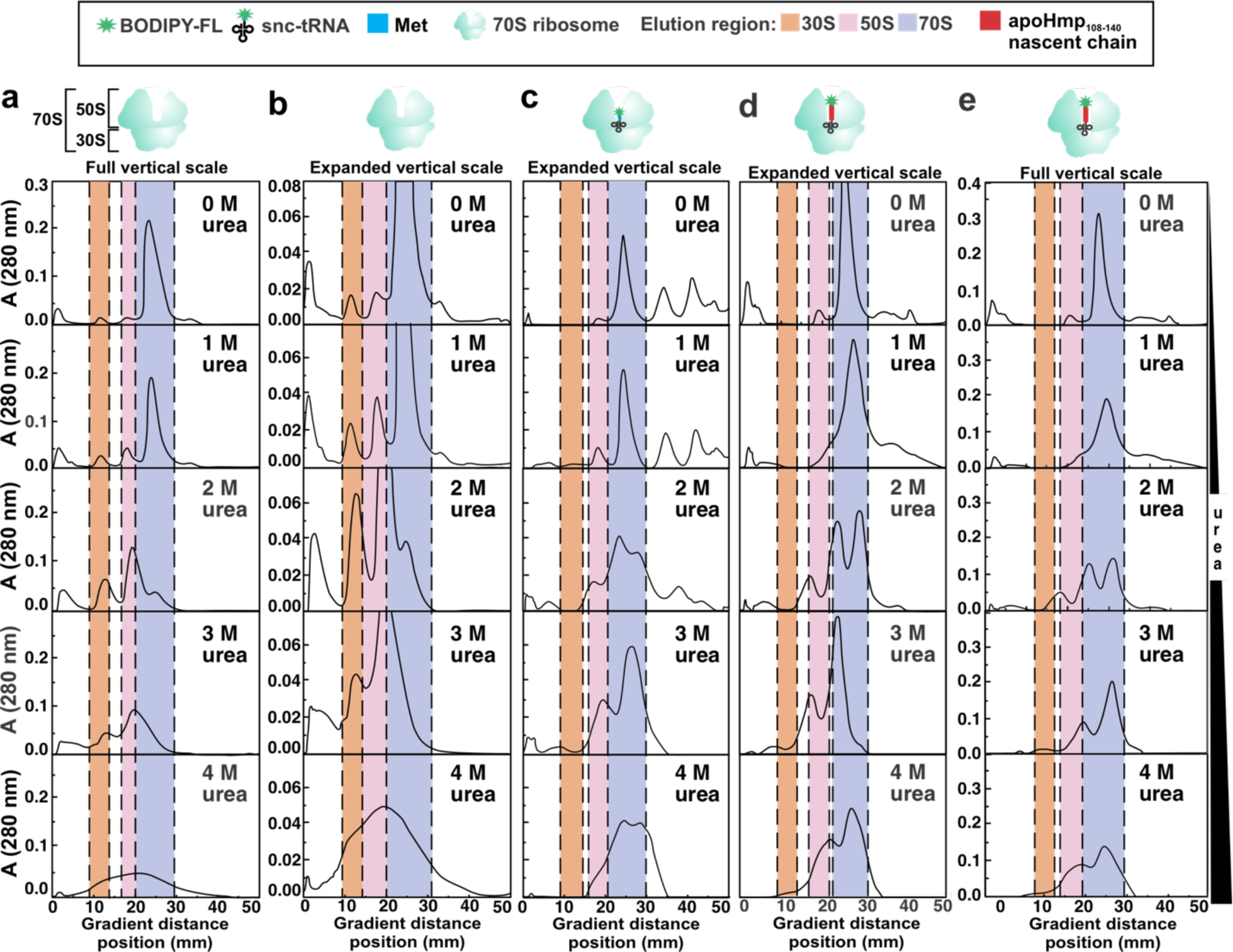
Sucrose gradient profiles for both empty 70S ribosomes (a), the same 70S ribosome profile is set to the same y-axis scale as the short (1-2 amino acid) nascent chains (b), and ribosomes harboring short (1-2 amino acid) nascent chains (c), ribosomes bearing a longer nascent-peptide chain (d), and the same ribosomes bearing a longer nascent-peptide chain at the full vertical scale (e) are shown for increasing concentrations of urea. Profiles were collected at 280 nm.

**Figure S4.**
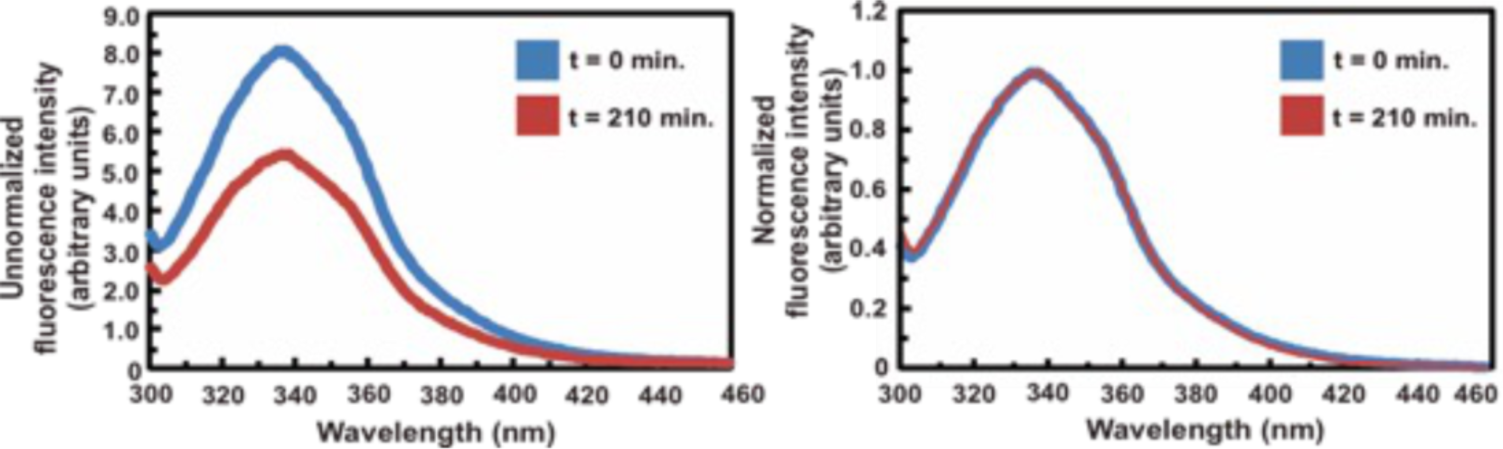
Time-course experiments for apoHmp_1-55_ are shown at 0.45 M urea. There is no change in spectral shift as incubation time is dramatically increased. Identification of ribosomal proteins crosslinked to apoHmp_1-140_ RNCs via Western blotting. Left, low-pH 10% SDS-PAGE analysis of N-terminal fluorescently-labeled apoHmp_1-189_ RNCs generated via transcription-translation in an *E. coli Dtig* S30 cell-free system followed by Western blotting (right) employing antibodies against ribosomal proteins (a) L24 and (b) L29. See Methods for Western blotting and antibodies details, representative data, out of n = 2, are displayed.

**Figure S5.**
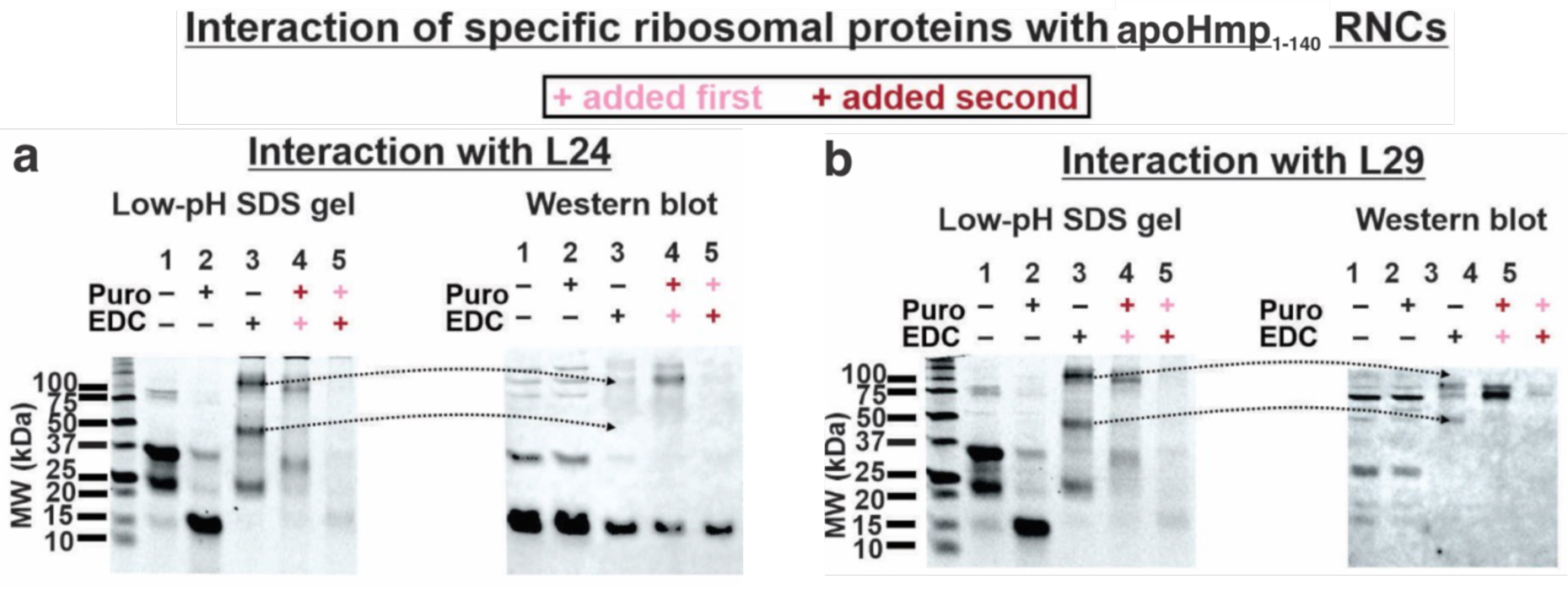
Identification of ribosomal proteins crosslinked to apoHmp_1-140_ RNCs via Western blotting. Left, low-pH 10% SDS-PAGE analysis of N-terminal fluorescently-labeled apoHmp_1-189_ RNCs generated via transcription-translation in an *E. coli Dtig* S30 cell-free system followed by Western blotting (right) employing antibodies against ribosomal proteins L24 and L29. See Methods for Western blotting and antibodies details, representative data, out of n = 2, are displayed.

**Figure S6.**
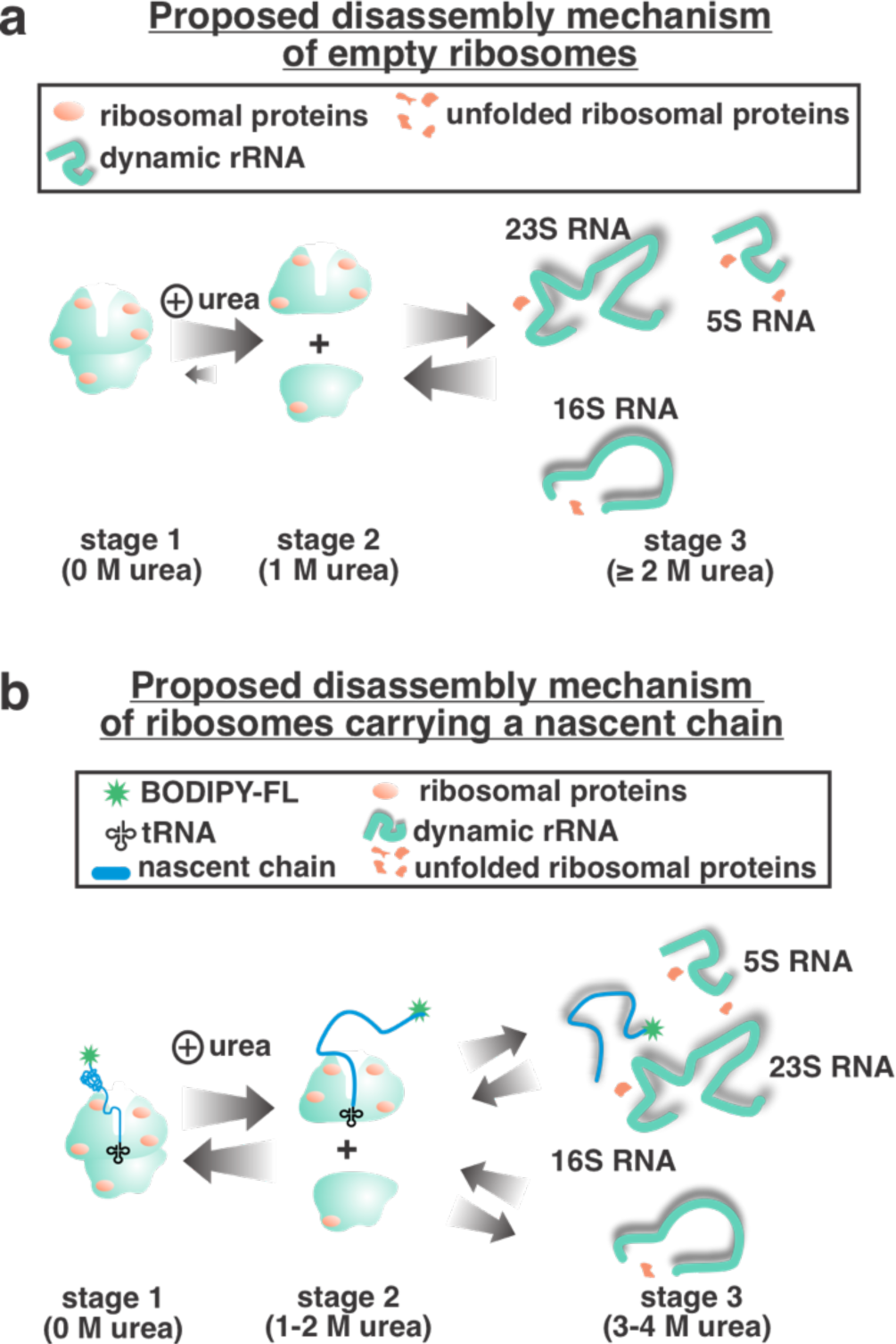
A proposed disassembly model for empty ribosomes and ribosome-bound nascent chains. (a) Briefly, empty ribosomes begin to dissociate from one another at c.a. 1 M urea. At > 2 M urea empty ribosomes experience r-protein and rRNA unfolding. (b) Briefly, RNCs begin to dissociate from one another at c.a. 1-2 M urea. At > 3 M urea empty ribosomes experience r-protein and rRNA unfolding.

**Figure S7.**
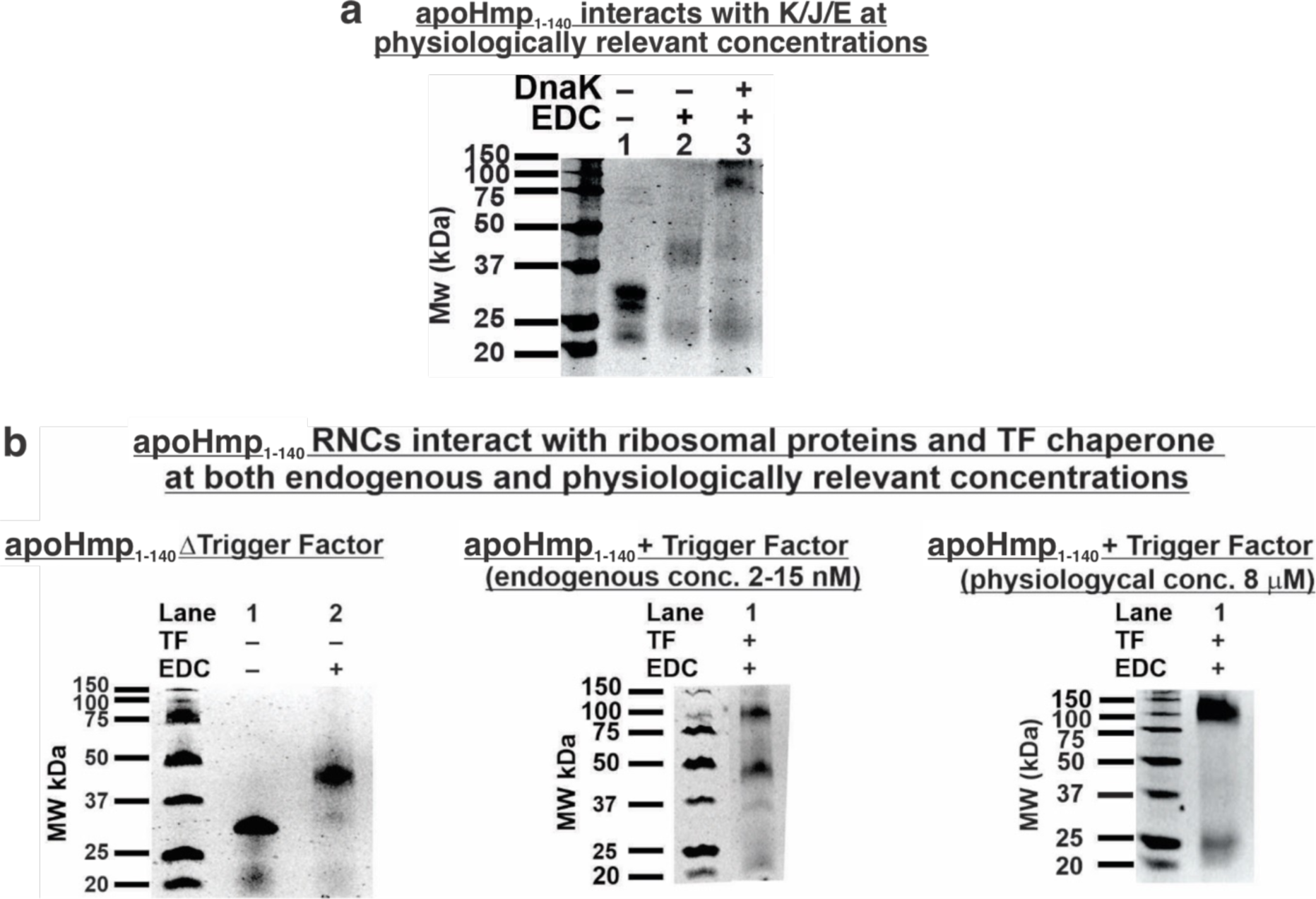
(a) Low pH 10% SDS-PAGE gel showing apoHmp_1-140_ with no Hsp70 chaperones added (K/J/E) and physiologically relevant concentrations, respectively. (b) ApoHmp_1-140_ at various concentrations of trigger factor molecular chaperone. Concentration of trigger factor increases from left gel to right gel. See methods for more information about chaperone concentration.

**Figure S8.**
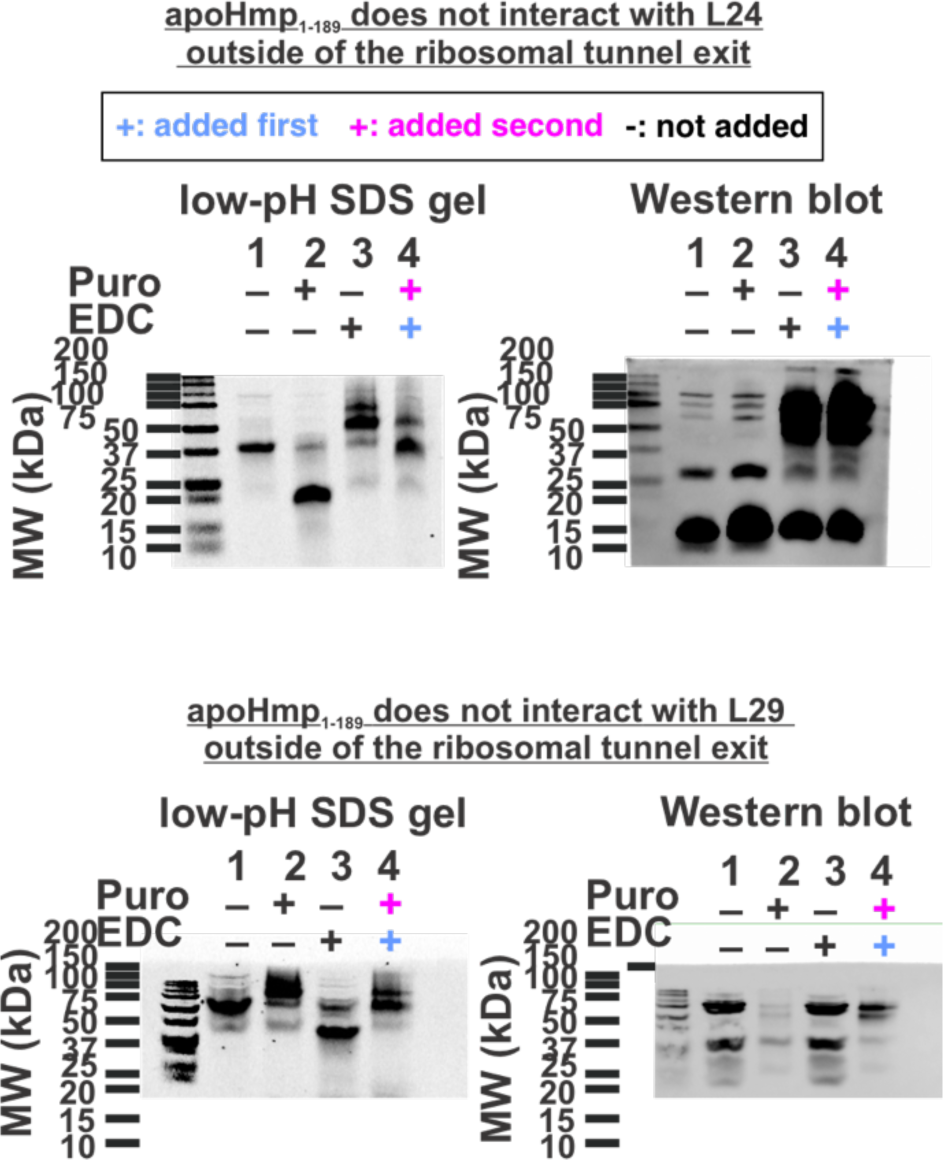
Identification of ribosomal proteins crosslinked to apoHmp_1-189_ RNCs via Western blotting. Left, low-pH 10% SDS-PAGE analysis of N-terminal fluorescently-labeled apoHmp_1-189_ RNCs generated via transcription-translation in an *E. coli Dtig* S30 cell-free system followed by Western blotting (right) employing antibodies against ribosomal proteins L24 and L29. See Methods for Western blotting and antibodies details, representative data, out of n = 2, are displayed.

